# Cytokine-induced reprogramming of human macrophages toward Alzheimer’s disease-relevant molecular and cellular phenotypes *in vitro*

**DOI:** 10.1101/2024.10.24.619910

**Authors:** Anna Podlesny-Drabiniok, Carmen Romero-Molina, Tulsi Patel, Wen Yi See, Yiyuan Liu, Edoardo Marcora, Alison M. Goate

## Abstract

Myeloid cells including brain-resident (microglia) and peripheral macrophages play a crucial role in various pathological conditions, including neurodegenerative disorders like Alzheimer’s disease (AD). They respond to disruption of tissue homeostasis associated with disease conditions by acquiring various transcriptional and functional states. Experimental investigation of these states is hampered by the lack of tools that enable accessible and robust reprogramming of human macrophages toward Alzheimer’s disease-relevant molecular and cellular phenotypes *in vitro*. In this study, we investigated the ability of a cytokine mix, including interleukin-4 (IL4), colony stimulating factor 1 (CSF1/MCSF), interleukin 34 (IL34) and transforming growth factor beta (TGFβ), to induce reprogramming of cultured human THP-1 macrophages. Our results indicate this treatment led to significant transcriptomic changes, driving THP-1 macrophages towards a transcriptional state reminiscent of disease-associated microglia (DAM) and lipid-associated macrophages (LAM) collectively referred to as DLAM. Transcriptome profiling revealed gene expression changes related to oxidative phosphorylation, lysosome function, and lipid metabolism. Single-cell RNA sequencing revealed an increased proportion of DLAM clusters in cytokine mix-treated THP-1 macrophages. Functional assays demonstrated alterations in cell motility, phagocytosis, lysosomal activity, and metabolic and energetic profiles. Our findings provide insights into the cytokine-mediated reprogramming of macrophages towards disease-relevant states, highlighting their role in neurodegenerative diseases and potential for therapeutic development.

## Introduction

Alzheimer’s disease (AD) is the most prevalent type of dementia in the elderly, affecting tens of millions of people worldwide; however, no effective disease-modifying treatments are currently available. Genome-wide association studies (GWAS) have identified genetic variants in ∼75 genomic regions (loci) associated with AD in people of European ancestry ^1^. Apolipoprotein E (*APOE)* is a major Alzheimer’s disease risk gene ^2^, with the *APOE* ε2 and ε4 alleles associated with reduced and increased risk of developing the disease respectively, compared to the most common *APOE* ε3 allele ^3^. We and others have shown that common non-coding AD risk alleles identified by GWAS are specifically enriched in enhancers that are active in macrophages including microglia, the resident macrophages and innate immune cells of the brain ^4–7^. These genetic and genomic findings strongly implicate macrophages in the etiology of AD. In addition, pathway analyses of AD GWAS data strongly implicate 1) cholesterol metabolism, 2) phagocytosis/endocytosis, and 3) the innate immune system in AD pathogenesis ^8,9^. These three biological pathways may not be independent causal drivers of AD; rather, they may be three components of a higher-order biological process acting as an AD pathogenetic hub in macrophages. This pathogenetic hub may be efferocytosis, one of the most fundamental activities of all macrophages, to find, phagocytose, digest and clear/dispose of apoptotic cells and other cholesterol-rich cellular waste ^10–12^. This multistep mechanism of efferocytosis is essential for the maintenance of tissue homeostasis and immune tolerance, and for the resolution of inflammation by eliminating damaged or otherwise unwanted cells and cellular debris (e.g., degenerating neurons, dystrophic neurites, myelin fragments and unwanted synapses) ^13^. When exposed to such debris, macrophages up-regulate the expression of several genes involved in phagocytosis, lysosomal processing, and cholesterol clearance. This gene expression profile is often referred to as DAM/LAM (for disease-associated microglia/lipid-associated macrophages) ^14–19^. As we proposed previously, we collectively refer to states of these microglia/peripheral macrophages as DLAM ^20^. The most up-regulated DLAM gene is *APOE*, a major gene for AD risk, cholesterol metabolism, and efferocytosis ^15,16,19,21,22^. Another AD risk gene, *TREM2*, was proposed to drive the transition from homeostatic to the DLAM state ^15,16^. Loss of TREM2 in mouse models and human iPSC-derived microglia hinders the ability of these cells to transition from a homeostatic state to the DLAM state, leading to impaired efferocytosis and response to amyloid plaques ^15,18,23–27^. In peripheral macrophages, loss of Trem2 also inhibits their transition to the LAM state, leading to adipocyte hypertrophy, systemic hypercholesterolemia, body fat accumulation, and glucose intolerance ^16^.

Recently, three approaches were proposed to recreate the DLAM state in human cell in vitro systems by 1) using lipid/cholesterol-rich efferocytic substrates like myelin debris, synaptosomes, and dead neuronal cells ^19^, 2) overexpressing a DLAM-inducing transcription factor ^19^ 3) reducing the expression of DLAM repressive transcription factors in iPSC-derived microglia ^20^. However, these iPSC-derived microglia models are expensive, not widely available and require engineered cell lines (knock-out or knock-in). Therefore, having a robust, widely accessible and reproducible *in vitro* system to modulate the expression of AD risk and DLAM genes that regulate efferocytosis and other disease-relevant cellular phenotypes may be a useful resource, particularly for functional genomic studies.

Here, we utilized an *in vitro* model of human immortalized THP-1 macrophages treated with a mix of anti-inflammatory cytokines such as IL4 and TGFβ, along with survival cytokines IL34 and MCSF, to polarize macrophages toward an anti-inflammatory, DLAM state ^28,29^). We observed a robust increased expression of DLAM genes, a decreased expression of proliferative genes and modulation of several AD risk genes when THP-1 macrophages were exposed to the cytokine mix as compared to control THP-1 macrophages. Importantly, unlike most previous studies, we performed extensive functional characterization of these DLAM-like macrophages *in vitro* probing each step of efferocytosis. This work offers insights into functional characteristics of THP-1 human macrophages with activated transcriptional programs similar to those observed in DLAM clusters.

## Results

### Cytokine mix treatment polarizes THP-1 macrophages to a DLAM-like transcriptional state

The main goal of this study was to develop an *in vitro* system for the investigation of Alzheimer’s disease-relevant molecular and cellular phenotypes in human macrophages. To this end we used the human THP-1 monocytic leukemia immortalized cell line that is more accessible, affordable, easier to manipulate when performing functional genomics experiments than microglia derived from induced pluripotent stem cells and can easily be differentiated to macrophages using phorbol 12-myristate 14-acetate (PMA). Differentiated cells were treated for 2-3 days with a combination of cytokines (cytokine mix) including IL4 (20ng/ml), MCSF (25ng/ml), IL34 (100ng/ml) and TGFβ (50ng/ml). We included IL4 and TGFβ to polarize cells toward an anti-inflammatory state. Previous studies showed that one of the DLAM regulators, LXRα, controls anti-inflammatory subsets of DLAM and promotes cholesterol clearance and immune response ^30^. Two additional cytokines, MCSF and IL34 were also selected to polarize human macrophages toward a more microglial-like phenotype by providing molecular cues present in the brain tissue microenvironment ^31,32^. In addition, IL4 and MCSF were previously shown to increase TREM2 expression ^28,29^, which is one of the key drivers of the DLAM state ^15,16^. Importantly, in human alveolar macrophages, IL4 has been shown to induce expression of well-known DLAM markers including CTSB, CTSD, LGALS3, APOE, APOC2 at the transcript and protein levels ^33^. As a control, we used THP-1 macrophages that were not exposed to the cytokine mix (Figure 1A). To assess global transcriptomic changes in THP-1 macrophages treated with the cytokine mix, we performed bulk RNA-seq. First, we used principal component analysis (PCA) to visualize the overall distance between the transcriptomes of control and cytokine mix-treated THP-1 macrophages and those of other myeloid cells and cell states publicly available from published studies. This analysis revealed that both control and cytokine mix-treated THP-1 macrophages, are separated from *ex-vivo* peripheral macrophages such as liver macrophages ^34^, macrophages from atherosclerotic plaques ^35^ and macrophages from adipose tissue ^16,35^. They show some degree of clustering with *ex-vivo* ^7,36,37^ and *in vitro* ^38^ primary microglia, but show a high degree of clustering with *in vitro* iPSC-derived microglia (iMGLs) ^19,20,34^ (Figure 1B). Next, we performed differential gene expression analysis (DGEA) and found 1,613 significantly up-regulated genes (logFC > 1, adj.p < 0.05), and 1,394 significantly down-regulated genes (logFC < 1, adj.p < 0.05). The top up-regulated genes include *AQP9*, *KRT16*, *CCL13*, *TGM2, CD274, SERPINB4, VCAM1*, while the most down-regulated genes include *GPR34, RGS18, COL21A1* (Figure 1D). Next, we used gene set enrichment analysis using curated canonical pathways such as KEGG, REACTOME (c2.kegg_legacy.v2023.2.Hs.symbols.gmt, c2.reactome.v2023.2.Hs.symbols.gmt) and Gene ontology gene sets collection (c5.mf.v2023.2.Hs.symbols.gmt, c5.bp.v2023.2.Hs.symbols.gmt, c5.cc.v2023.2.Hs.symbols.gmt) to identify changes in THP-1 macrophages driven by cytokine mix treatment. We found “Oxidative phosphorylation” (M19540), “Lysosome” (M11266), “Retinol Metabolism” (M9488), “PPAR signaling pathway” (M13088), “Fatty Acid metabolism (M699) among the top positively enriched KEGG pathways (Figure 1C), while, “FC Gamma Receptor Mediated Phagocytosis” (M16121), “Gap junctions” (M4013), “Ribosome” (M189), and “Cell Cycle” (M7963) were among the top negatively enriched KEGG pathways (Figure 1C). To investigate whether changes in gene expression and pathways in cytokine mix treatment are driven by IL4 or three factors (3F) of survival/maturation cytokines (MCSF, IL34, TGFβ) we conducted bulk RNAseq and gene expression analysis using THP-1 macrophages treated with IL4 only and 3F only and compared them to the control condition. We found that IL4 is the main driver of differential gene expression and pathway enrichment in cytokine mix-treated THP-1 macrophages as evidenced by similar top up- and down-regulated genes and normalized enrichment score of top canonical pathways changed in cytokine mix-treated THP-1 macrophages (Figure 1C-D). Results of differential gene expression analysis and gene set enrichment analyses for all cytokine treatment groups including single and cytokine mix-treatments as compared to non-treated THP-1 macrophages are included in Table S1.

**Figure 1.**
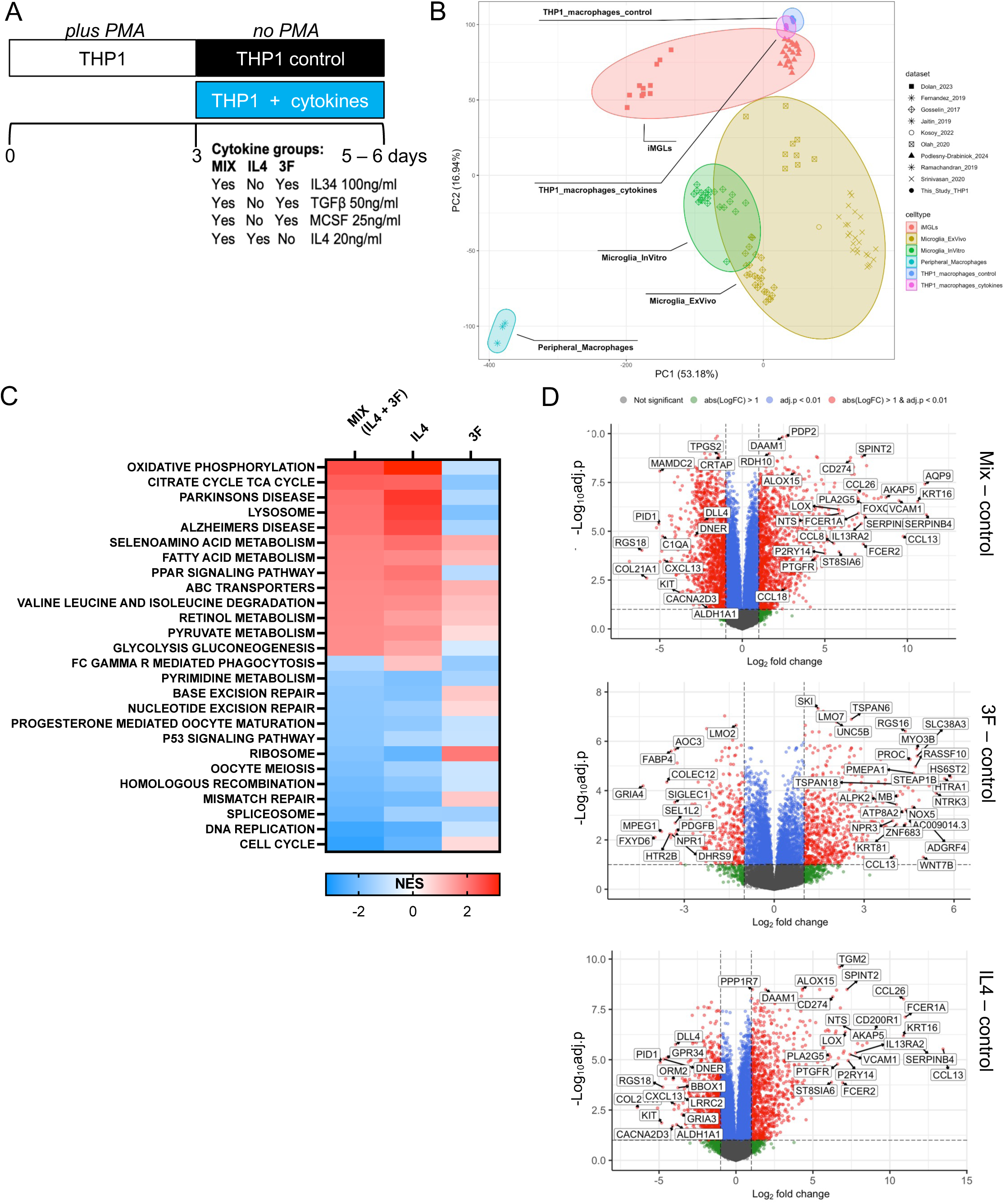
Differential abundance and pathway enrichment analysis of gene expression changes induced by cytokine treatment in THP-1 macrophages reveales changes in Lysosome, Oxidative phosphorylation and Cell cycle. A) Schematic of THP-1 differentiation using phorbol 12-myristate 13-acetate (PMA) followed by cytokine treatment B) PCA analysis comparing THP1 macrophages (with and without cytokines) with other myeloid cell types C) KEGG-selected pathways from Gene set enrichment analysis (GSEA) positively in three main cytokines group as compared to No cytokine (control) (for all GSEA see Table S1) D) Volcano plot representing top differentially expressed genes in cytokine-treated macrophages.

Next, we used gene set enrichment analysis (GSEA) to test whether our cytokine mix treatment effectively induced transcriptional DLAM response in THP-1 macrophages. To this end, we used the following DLAM gene sets: 1) humanized DAM up- and down-regulated gene sets from mouse models of amyloidosis ^15,39–43^ 2) DAM-like gene sets from human AD brains ^36,44–47^, 3) LAM that reside in a lipid-rich niche ^16,35,48,49^, 4) *in vitro* DAM obtained by exposure to lipid-rich phagocytic substrates ^19^, 5) Other gene sets including Proliferative, Interferon and LPS gene sets ^42,45,47^. All gene sets are listed in Table S2 and have a unique gene set identifier (GS) used throughout this manuscript. Overall, we found that transcriptional changes that occur in cytokine mix-treated THP-1 macrophages are positively enriched for genes up-regulated in disease-associated microglia clusters such as DAM ^15^ (GS1) and ARM ^41^ (GS5), and microglial signatures from Neurodegeneration module ^42^ (GS10), CD11c positive microglia ^40^ (GS3), and Aβ-associated microglia ^39^ (GS7) that were selected from mouse models of amyloidosis (Figure 2A). Importantly, we showed positive enrichment of genes up-regulated and negative enrichment of genes down-regulated in clusters closely resembling mouse DAM obtained from sc/sn-RNAseq of microglia from AD brains in cytokine mix-treated THP-1 macrophages compared to control THP-1 macrophages. We found positive enrichment of the AD1 signature from Aβ-associated microglia ^44^ (GS11), Cluster 7 up-regulated genes that mostly resembled mouse DAM ^36^ (GS19), lipid-processing microglia (MG4), and glycolytic microglia (MG7) signatures ^47^ (GS21 and GS22), GPNMB_NACA, GPNMB_EYA2 and LPL_CD83 DAM cluster signatures from AD brain biopsies ^45^ (GS13-GS15) in cytokine mix-treated THP-1 macrophages (Figure 2A). In addition, we found that transcriptional changes that occur in cytokine mix-treated THP-1 macrophages are positively and negatively enriched for, respectively, genes up- and down-regulated in Cluster 2 and 8 of human iPSC-derived microglia treated with lipid-rich brain phagocytic substrates *in vitro* ^19^ (GS29-32). Similar transcriptomic activation states to those identified in microglia in aging and AD brains have been observed in subpopulations of macrophages in diseased lipid-rich tissues (e.g., TREM2^high^ macrophages in atherosclerotic plaques ^49^, lipid-associated macrophages (LAM) in fatty liver and obese adipose tissue ^16^). Therefore, we examined the enrichment of signatures from lipid-associated macrophages in THP-1 macrophages treated with cytokine mix compared to control. We found statistically significant, positive enrichment of genes up-regulated in LAM from obese individuals ^16^ (GS25), foamy macrophages ^35^ (GS24), humanized signature from TREM2^high^ macrophages from atherosclerotic plaques ^49^ (GS23) and white matter microglia isolated from patients with multiple sclerosis that are exposed to large demyelinated lesions ^48^ (GS27) with concomitant negative enrichment of genes down-regulated in these clusters (i.e. genes down-regulated in LAM ^16^ (GS26) and in white matter microglia from MS patients ^48^ (GS28)) (Figure 2A). Consistent with the GSEA results, we also found an increased expression of selected DLAM genes in cytokine mix-treated macrophages in bulk transcriptomes (Figure 2B).

**Figure 2.**
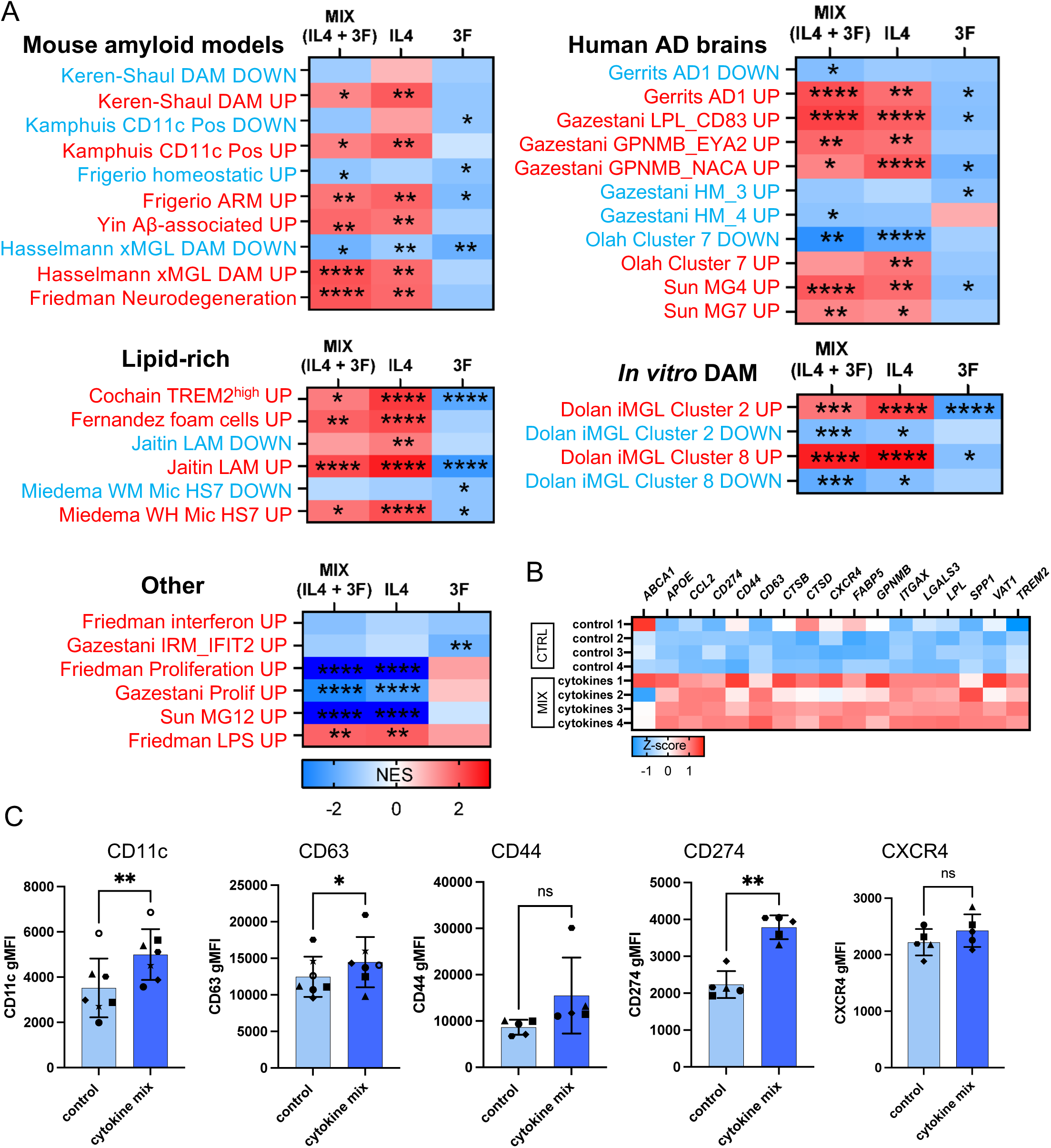
Cytokine treatment induces a DLAM, and non-proliferative transcriptional response in THP-1 macrophages. A) Enrichment of AD-related gene sets in Cytokine-treated macrophages. All gene sets are listed in Table S2, NES – normalized enrichment score * FDR q-val < 0.05 ** FDR q-val < 0.01 *** FDR q-val < 0.001 **** FDR q-val < 0.0001. B) Heatmap of DLAM genes Z-scores (log2 transformed TPM) based on Table S1 C) Quantification of surface expression of DLAM markers by flow cytometry. Values plotted as gMFI. Groups were tested with paired t.test (two-tailed) * p < 0.05, ** p < 0.01, *** p < 0.001. Percentage of positive cells plotted separately in Figure S2. Different dot shapes correspond to independent macrophage differentiations. N = 3-4 independent differentiation, each differentiation has 2 well replicates.

Apart from DLAM subsets, several other subsets of microglia/peripheral macrophages have been identified by scRNAseq including subsets involved in Proliferation, Interferon response, LPS-induced, which were examined by GSEA. We found a strong, negative enrichment of genes up-regulated in the Proliferative cluster (Prolif) ^45^ (GS33) and cycling microglia (MG12) ^47^ (GS34) found in AD brains as well as negative enrichment of a humanized Proliferation module^42^ (GS36) (Figure 2A). This is also in agreement with gene set enrichment analysis showing “Cell cycle” as the top negatively enriched pathway (Figure 1C). Intriguingly, we found positive enrichment of the LPS-induced module ^42^ (GS37) in cytokine mix-treated macrophages (Figure 2A). Additionally, some of the LPS-induced genes such as *CCL2* (Figure 2B) *CD274*, and proteins CD44, CD274 (Figure 2C) were markedly increased. Indeed, a number of studies show IL4 promotes expression of both anti- and pro-inflammatory genes in human monocytes and macrophages ^50,51^ and importantly, DLAM has been shown to be a heterogenous population enriched for both pro- and anti-inflammatory subsets ^30^.

To dissect which cytokine is driving the effect of DLAM polarization and negative impact on Proliferative clusters, we included two additional conditions: single IL4 treatment and 3F and performed GSEA analysis. Similarly to global transcriptomic changes, we demonstrated that positive enrichment of DLAM gene sets and negative enrichment of homeostatic and proliferative gene sets is driven by IL4 rather than 3F as evidenced by very similar values of NES between cytokine mix and IL4 (Figure 2A).

To further validate transcriptomic findings and the positive enrichment of DLAM markers in THP-1 macrophages treated with our cytokine mix, we performed flow cytometric studies to assess the level of surface DLAM markers. We used previously validated classic DLAM markers such as CD11c ^15,30,44^ and CD63 ^16^, CXCR4, a marker that has previously been shown to be an anti-inflammatory DLAM surface marker, and two pro-inflammatory DLAM markers CD44 and CD274 ^30^. Consistent with transcriptomic changes, we found statistically significant increased DLAM geometric fluorescent intensity (gMFI) of CD11c, CD63, and CD274 in THP-1 macrophages treated with the cytokine mix as compared to control macrophages (Cytokine mix - Control Mean of differences [95% confidence interval], p=two-tailed paired t test gMFI (CD11c: 1473 [862.9 to 2083] p=0.001, CD63: 1996 [183.9 to 3809] p=0.035, CXCR4: 206.3 [−206.3 to 618.8] p=0.237, CD44: 6863 [−4701 to 18427] p=0.174, CD274: 1553 [1043 to 2063] p=0.001) (Figure 2C, Figure S2). We did not observe an increased percentage of positive cells suggesting a higher level of DLAM marker gene expression per cell rather than increased proportion of DLAM-like cell population (Figure S2) (% of positive cells (CD11c: 3.564 [−12.17 to 19.30] p=0.5996, CD63: −3.39 (−8.56 to 1.77) p=0.158, CXCR4: 1.99 [−0.29 to 4.28] p=0.017, CD44: −2.01 [−7.99 to 3.97] p=0.403, CD274: 42.88 [32.40 to 53.36] p<0.001]). One exception to that was CD274, previously associated with the pro-inflammatory subset of DLAM ^30^. It was weakly expressed on the surface of control cells but induced in cytokine mix-treated THP-1 macrophages (Figure S1). This is also in agreement with the transcriptomic level of *CD274*, which was one of the top up-regulated genes in cytokine mix-treated THP-1 macrophages (Figure 1D). Altogether, these data show that the mix of cytokines including anti-inflammatory and microglial maturation cytokines promotes DLAM-like transcriptional responses in THP-1 macrophages that recapitulate both anti- and pro-inflammatory transcriptional signatures identified by weighted co-expression network analysis ^30^.

### Candidate AD risk genes are differentially expressed in cytokine mix-treated THP-1 macrophages

Multiple integrative analyses of AD GWAS and functional genomic data have identified candidate AD risk genes expressed in myeloid cells ^4–7^. Some of the candidate AD risk genes are upregulated in DLAM states including *APOE* and *TREM2*, the cholesterol transporter *ABCA1* and lysosomal markers such as *CTSB*, and *GRN*. Similarly, several AD candidate genes are downregulated in DLAM states including *MS4A6A*, *BIN1*, *CD2AP*, and *RIN3* suggesting that AD risk variants may impact the plasticity of microglia to transition from homeostatic to DLAM state. Using a list of 81 candidate AD risk genes nominated by ^1,52^ we found that more than half of them (44/81) are differentially expressed in cytokine mix-treated THP-1 macrophages (adj.p < 0.05) including *PICALM, SCIMP, PTK2B, FERMT2, ABI3, HLA-DQA1, APOE* (Figure S1A). This effect is mainly driven by IL4 which, when applied alone, induces differential expression of 36/81 genes. 3F alone induced differential expression of one fourth of AD risk genes (21/81). Five genes (5/81) were excluded because they did not appear in the bulk transcriptome or were filtered out because of very low expression level (*CLNK*, *PRDM7*, *IL34*, *HS3S35T*, *TSPOAP1*). The list of candidate AD risk genes and their expression level in cytokine-treated groups as compared to control is listed in Table S2.

Next, we used a list of genes whose expression levels in myeloid cells are predicted to alter disease susceptibility, from an integrative analysis of AD GWAS and myeloid genomic data that we previously conducted ^6^. In Novikova et al. we found 29 candidate causal genes whose directionality of expression (increased or decreased level) was predicted to increase AD risk (see Fig.5 in ^6^). We added *TREM2* and *APOE* to this list, since their directionality of expression has been previously associated with modulation of AD risk ^53–55^. In particular the lower expression level of APOE has been genetically associated with higher risk of AD ^55^. The list of 31 myeloid candidate AD risk genes is depicted in Figure S1, and listed in Table S2. As proposed by Novikova et al., ^6^, red indicates that increased expression of the gene is predicted to increase risk for AD, blue indicates that decreased expression of the gene is predicted to increase risk for AD, gold indicates that the directionality of gene expression that is associated with increased disease susceptibility cannot be robustly inferred. We filtered log2 fold change (logFC) for each gene in three conditions (cytokine mix, IL4 and 3F) and attributed change in a contrast (cytokine(s) treatment vs control) to increased or decreased AD risk. For example, decreased expression of *BIN1* was associated with increased AD risk ^6^. In cytokine mix-treated THP-1 macrophages, *BIN1* level was significantly increased as compared to non-treated macrophages (logFC = 1.68 adj.p = 1.04e-5), therefore the effect of cytokine mix treatment on *BIN1* expression in THP-1 macrophages is indicative of a risk-decreasing effect and is labeled as “risk-decreasing” in the heatmap (Figure S1B). On the contrary, *BIN1* expression is significantly reduced upon exposure to 3F only (logFC = −0.83, adj.p = 0.019) as compared to non-treated THP-1 macrophages, suggesting 3F treatment in THP-1 macrophages modulates *BIN1* expression toward increased AD risk and was therefore labeled as “risk-increasing” in the heatmap (Figure S1B). The level of *BIN1* expression in IL4-treated THP-1 macrophages was not significantly changed as compared to non-treated macrophages (logFC = 0.30, adj.p = 0.26), therefore we cannot reliably infer the effect on AD risk and (no label on the heatmap) (Figure S1B).

We observed a group of genes strongly implicated in efferocytosis ^12^ such as *AP4M1, ZYX, PTK2B, PILRA, BIN1, APOE* and *TREM2* whose increased levels of expression in cytokine mix treated THP-1 macrophages are predicted to reduce AD risk. Another interesting observation was that IL4 and 3F have different effects on *MS4A4A* and *MS4A6A*, whose increased levels of expression are predicted to increase AD risk ^6^. We found that in THP-1 macrophages the expression levels of these two genes are induced by IL4 treatment while they are significantly reduced by 3F treatment. Interestingly, in cytokine mix-treated THP-1 macrophages, the effect of 3F dominated that of IL4 leading to decreased expression of *MS4A4A* and *MS4A6A*, which is in turn associated with reduced AD risk (Figure S1B). Finally, in cytokine mix-treated THP-1 macrophages, there was a group of genes (including *SPI1*, *SCIMP*, and *REBEP1*) whose increased expression levels are associated with increased AD risk.

In summary, cytokine mix treatment induced the expression of ten myeloid candidate AD risk genes toward decreased risk and modulated the expression of five other myeloid candidate AD risk genes toward increased risk (Figure S1B). IL4 treatment induced expression of six myeloid candidate AD risk genes toward protection and nine genes toward increased risk. Finally, 3F induced the expression of five myeloid candidate AD risk genes toward protection and four genes toward increased risk (Figure S1B).

Altogether we found that the cytokine mix treatment which includes a combination of IL4, and 3F (MCSF, IL34 and MCSF) is the most effective treatment compared to single 3F or IL4 in 1) induction of AD risk gene expression 2) induction of myeloid candidate AD risk genes in the same direction as associated with decreased AD risk. Hence, we decided to conduct further analysis including single cell RNAseq and functional studies using cytokine mix treatment and control non-treated THP-1 macrophages (control) excluding 3F and IL4 conditions.

### Single-cell RNA-seq identifies disease-associated and non-proliferative transcriptional states in THP-1 macrophages treated with cytokine mix

To confirm our bulk RNA-seq findings and test if exposure to the cytokine mix induced different transcriptional states in THP-1 macrophages cultured *in vitro*, we performed scRNA-seq on control and cytokine mix-treated cells. Quality control, integration and clustering were performed prior to downsampling cells to the smallest treatment condition for comparison, resulting in 27,584 total cells (Figure 3A; Table S3). Cluster annotations were defined based on the enrichment of existing myeloid gene expression signatures and biological pathways (Figure 3B; Figure S4A). We found nine unique clusters Mac0-Mac8 where Mac0 showed increased expression of DLAM canonical markers such as *CTSB*, *CTSZ*, *SPP1*, *ITGAX* and *CLEC7A* (Figure 3B, Figure S3A,B). We employed several methods to compare our data with existing datasets from brain-resident and peripheral macrophages from *in vitro* and *in vivo* studies that identified different macrophage transcriptional states. First, we tested which THP-1 macrophage clusters overlap with published DLAM cluster marker genes. We found that the cluster marker genes identified in this study showed a statistically significant hypergeometric overlap with DLAM cluster marker genes including marker genes from AD brains such as GPNMB_NACA cluster (GS15) (hypergeometric overlap adjusted p value: adj.p_Mac0_ = 5.90e-17, adj.p_Mac3_ = 5.60e-19, adj.p_Mac4_ = 0.075, adj.p_Mac7_ = 0.009, adj.p_Mac8_ = 1.80e-13), GPNMB_EYA2 cluster (GS14) (adj.p_Mac0_ = 1.80e-09, adj.p_Mac4_ = 0.017, adj.p_Mac7_ = 1.00e-61, adj.p_Mac8_ = 0.016), LPL_CD83 cluster (GS13) (adj.p_Mac0_ = 6.40e-09, adj.p_Mac3_ = 0.050, adj.p_Mac4_ = 0.030, adj.p_Mac7_ = 8.60e-26, adj.p_Mac8_ = 4.90e-06) identified by Gazestani et al. ^45^, MG4 (lipid processing) cluster (GS21) described by Sun et al. (adj.p_Mac0_ = 1.00e-06, adj.p_Mac4_ = 0.001, adj.p_Mac7_ = 9.80e-53) ^47^, LAM signature (GS25) from peripheral macrophages from obese individuals ^16^ (adj.p_Mac0_ = 8.20e-10, adj.p_Mac3_ = 2.90e-11, adj.p_Mac4_ = 0.006, adj.p_Mac8_ = 3.50e-26), and cluster marker genes from white matter microglia isolated from patients with multiple sclerosis (HS7 cluster) (GS27) ^48^ (adj.p_Mac0_ = 4.80e-08, adj.p_Mac3_ = 1.20e-37, adj.p_Mac8_ = 2.20e-22). In addition we found that marker genes from Mac0, Mac3, Mac4, Mac7, Mac8 showed overlap with *in vitro* DAM cluster marker genes such as Cluster 2 (GS29) (adj.p_Mac0_ = 4.30e-29, adj.p_Mac3_ = 0.029, adj.p_Mac4_ = 3.60e-04, adj.p_Mac7_ = 1.80e-12, adj.p_Mac8_ = 4.00e-12) and Cluster 8 (GS31) (adj.p_Mac0_ = 5.40e-27, adj.p_Mac3_ = 7.10e-04, adj.p_Mac4_ = 4.80e-05, adj.p_Mac7_ = 1.50e-04, adj.p_Mac8_ = 1.60e-14) identified by Dolan et al, ^19,45^, marker genes from DAM cluster extracted from 5xFAD brain (GS1) (adj.p_Mac0_ = 4.90e-11, adj.p_Mac3_ = 1.20e-17, adj.p_Mac4_ = 0.002, adj.p_Mac7_ = 0.220, adj.p_Mac8_ = 4.80e-14) ^15^, as well as, marker genes from DAM cluster from xenografted human iPSC-derived microglia from 5xFAD mouse (GS8) (adj.p_Mac0_ = 4.80e-16, adj.p_Mac3_ = 5.50e-12, adj.p_Mac4_ = 0.007, adj.p_Mac7_ = 9.40e-5, adj.p_Mac8_ = 1.90e-18) ^43^ (Figure 4A). These results show that we did not identify one DLAM cluster in THP-1 macrophages, instead we found increased enrichment of DLAM genes in multiple clusters (Mac0, Mac3, Mac4, Mac7, Mac8) (Figure S3A,B; Table S3). Mac1 showed a strong hypergeometric overlap with marker genes extracted from Proliferative clusters identified by others in AD brains, Gazestani Cluster Prolif (GS33) (adj.p_Mac1_ = 2.40e-31) ^45^, Sun MG12 (cycling microglia) (GS34) (adj.p_Mac1_ = 5.80e-239) ^47^. Next, we tested the enrichment of the DLAM cluster marker genes in the THP-1 dataset calculating module score for several DLAM transcriptional signatures. We found that Mac0 and Mac8 showed the highest enrichment of Cluster 2 (GS29) and Cluster 8 (GS31) marker genes ^19^ (Figure 4C). Cluster 2 and 8 appear in iMGLs upon exposure to brain phagocytic substrates such as apoptotic neurons (Cluster 2 and 8), Aβ fibrils (Cluster 8), and myelin fragments and synaptosomes (Cluster 2) ^19^. Interestingly, we have also found an enrichment of DLAM signatures from AD brains in Mac0 and Mac6 clusters. Lipid processing cluster marker genes (MG4) (GS21) from AD brains identified by ^19,47^ and cluster marker genes from GPNMB_EYA2 (GS14) and LPL_CD83 (GS13) clusters identified in AD biopsies ^45^ were particularly enriched in Mac6 and part of Mac0 cluster identified in this study (Figure 4C, Figure S3C. This observation that microglial state found in AD brains exists in a small population of human immortalized macrophages cultured in a dish is very intriguing and underscore the utility of *in vitro* models to recapitulate some of the states found in the human brain. In addition, cluster marker genes of the My2 macrophage cluster (GS51) isolated from human atherosclerotic plaques showed enrichment with Mac0, Mac8, and Mac3 clusters ^56^ (Figure S3C) emphasizing the shared responses of peripheral and brain-resident macrophages to a lipid-rich environment. Finally, we found that proliferative states that exist in *in vitro* iMGLs exposed to brain phagocytic substrates ^19^ and human AD brains ^19,45^ are highly enriched in Mac1 cluster that was also annotated in this study as Proliferative and showed a high degree of overlap with known proliferative cluster marker genes (Figure 4A,C). Intriguingly, when we tested for enrichment of AD risk genes (Table S2) nominated as candidate causal genes in AD GWAS loci ^1,12^, we found an enrichment of AD risk genes in DLAM clusters such as Mac0, Mac2, Mac8, and additionally in Mac6 which is the most relevant to human DAM and lipid processing DLAM clusters (Figure 3C, Figure S3C).

**Figure 3.**
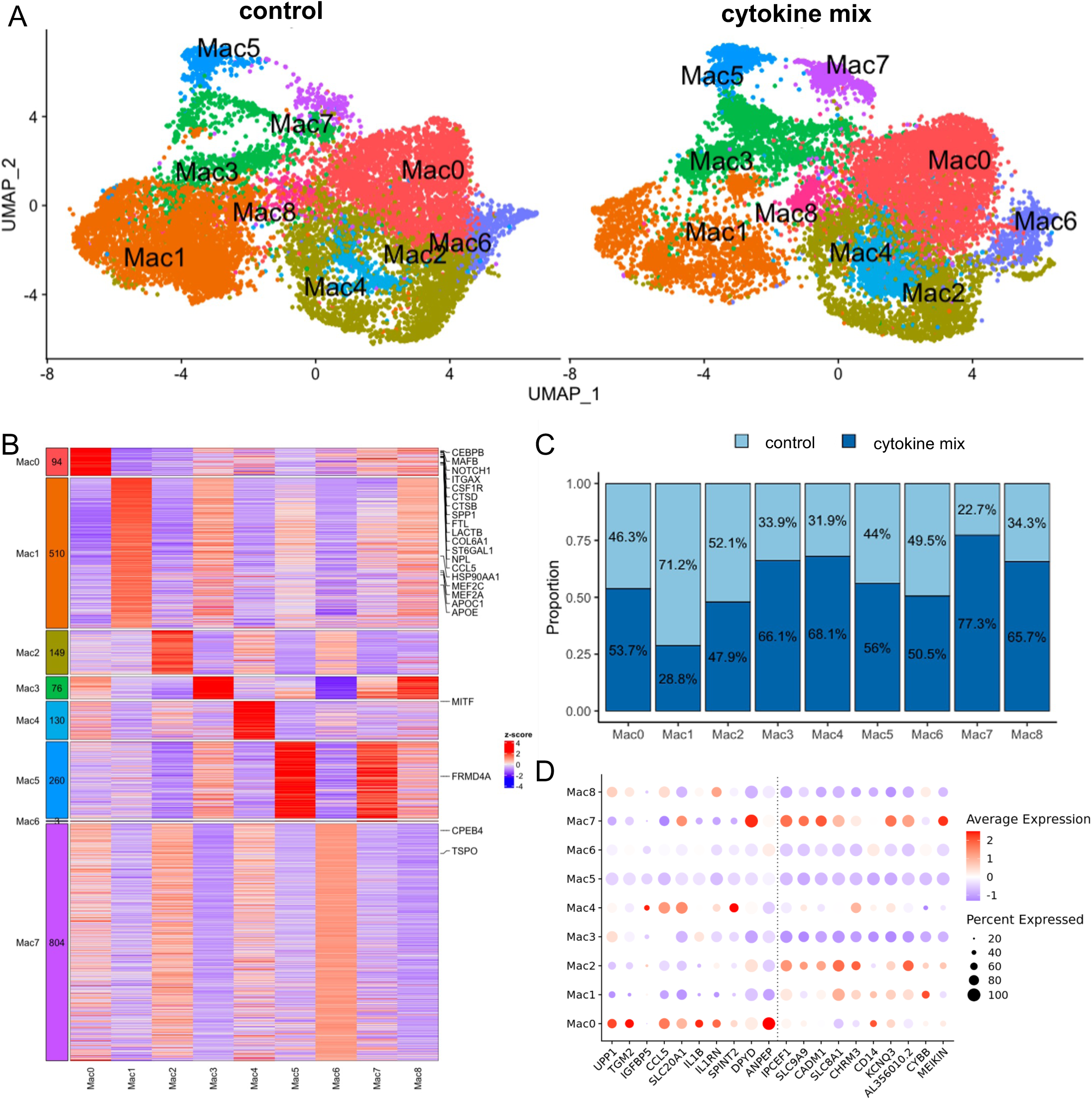
Cytokine mix treatment induces DLAM, non-proliferative macrophage transcriptional states in THP-1 macrophages. A) UMAP of downsampled scRNA-seq data comparing THP1 macrophages at baseline (left) and following cytokine treatment (right). Nine independent clusters were identified using Seurat FindClusters. B) Heatmap showing top expressed genes per cluster compared to all other clusters (log2FC>0.6, qval<0.05, pct expressed>70%). The left block shows the number of genes in each geneset and representative genes are labeled on the right. Z-scores across clusters are used for the plot. C) The proportion of cells per treatment condition within each cluster after downsampling. The percentage of cells from the total number of cells per cluster is shown on each bar. D) Dotplot showing expression of the top 10 differentially expressed genes across each cluster. The size of dots are scaled to represent the percentage of cells expressing the gene within the cluster. Average expression is normalized within each feature.

**Figure 4.**
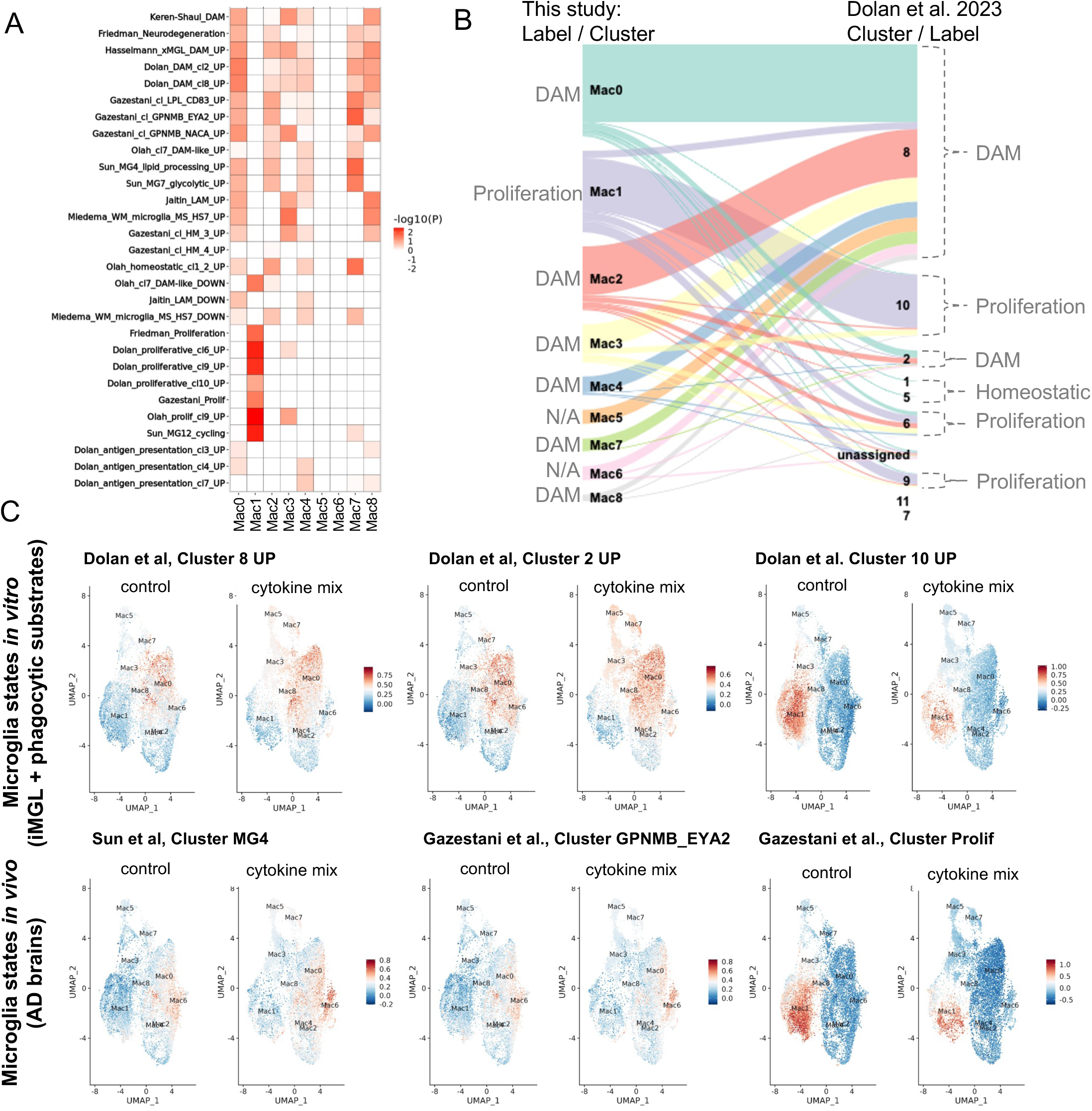
Comparison of THP-1 macrophages treated with cytokine mix with iPSC-derived microglia (iMGLs) exposed to phagocytic substrates and AD brain signatures reveals induction of similar transcriptional states. A) Hypergeometric overlap results showing enrichment of myeloid gene signatures in the up-regulated differentially expressed genes from pseudobulk scRNA-seq of cytokine treatment vs controls, grouped by subtypes. B) Sankey plot showing the projection of clusters found in this study to clusters identified by Dolan et al., by exposure of iMGLs to phagocytic substrates. C) UMAP projection of THP-1 macrophages dataset, cells colored by module scores of transcriptional signatures identified in DAM Clusters 2 and 8, and Proliferative Cluster 10 by Dolan et al., 2023; Lipid processing Cluster MG4 in AD brains identified by Sun et al, 2024, and DAM cluster GPNMB_EYA2 and Prolif cluster in AD biopsies identified by Gazestani et al., 2024.

Next, we compared the THP-1 transcriptional states with those found *in vitro* in iMGLs exposed to brain phagocytic substrates ^19^. We projected our THP-1 macrophage clusters into clusters identified by Dolan et al, and found that the majority of our macrophage clusters project to the DAM Cluster 8 (Figure 4B). This is also in agreement with our cluster annotation and hypergeometric overlap that show that multiple macrophage clusters identified in this study showed increased expression of DLAM genes. Consistently, all cells from Mac0 were projecting to Cluster 8 confirming Mac0 is the most representative of our DAM clusters (Figure 4B). We have also confirmed the existence of Proliferative clusters in THP-1 macrophages and found that Mac1 Proliferative cluster cells project to three proliferative clusters identified in iMGLs (Cluster 10, 6, 9) (Figure 4B). Next, we separately projected clusters from control conditions only (“control”) and from treatment condition only (“Cytokine mix-treated”) and we found that in control, cells from cluster Mac0 are distributed among Dolan’s Cluster 8 (n=2,476 cells) and 2 (n=601 cells), while in cytokine mix-treated macrophages nearly all cells from Mac0 (n=3,786 cells) projected to Cluster 8 (Figure S3D). We have also found that many more cells in control THP-1 macrophages projected to Proliferative clusters (Cluster 10, 9 or 6) identified by Dolan et al., (n=5,026 cells), whereas in cytokine mix-treated THP-1 cells only n=2,360 cells were found in the Proliferative clusters from Dolan et al (Figure S3D) confirming depletion of proliferative and induction of DLAM states in cytokine mix-treated macrophages.

To find biological processes enriched in each cluster we performed gene set enrichment analysis using gene ontology terms (GO) and cluster marker genes for each cluster. Mac0 cluster marker genes were enriched in “Regulation of Phagocytosis” (GO:0050764), “Receptor-mediated Endocytosis” (GO:0006898), “Intracellular pH reduction” (GO:0051452) (Figure S4A). In addition to that, CLEC7A, the canonical DAM marker associated with microglial response to amyloid plaques and apoptotic neurons ^17,57^ was enriched in Mac0 cluster (Figure S3B). Genes from clusters Mac3 and Mac8 were enriched in biological processes involved in “Cytoplasmic translation” (GO:0002181), “Peptide Biosynthetic Process” (GO:0043043), and “Ribosome Biogenesis” (GO:0042254) (Figure S4A). Mac4 genes were enriched for “Regulation of Neuron death” (GO:1901214) and “Leukocyte Tethering or Rolling” (GO:0050901). (Figure S4A). Biological processes enriched in genes positive in cluster Mac7 were “Regulation of Transcription by RNA Polymerase” (GO:0006357), “Circulatory System Development” (GO:0072359), “Positive Regulation of GTPase Activity” (GO:0043547). Mac1 genes were enriched for biological processes involved in “Mitotic Sister Chromatid Segregation” (GO:0000070), “DNA Metabolic Process” (GO:0006259), “Positive Regulation of Cell Cycle Process” (GO:0090068) (Figure S4A).

To determine whether cytokine mix exposure had any significant effects on the proportion of THP-1 macrophage subtypes, we used the *propeller* method ^58^ with adjustment for two independent macrophage differentiations. We observed an increase in the proportions of several DLAM-like clusters in response to cytokine mix treatment including Mac0 (29.1% - 24.7% = 4.4% [3.9%, 4.4%] cytokine mix-treated - control, t-statistics = −0.46, p = 0.647), Mac3 (14.8% - 7.1% = 7.7% [7.1%, 8.4%], t-statistics = −1.73, p = 0.099), Mac4 (7.5% - 3.3% = 4.2% [3.5%, 4.9%], t-statistics = −1.81, p = 0.086), Mac7 (6.2% - 1.6% = 4.6% [3.3%, 5.9%], t.statistics = −2.80, p = 0.011), and Mac8 (3.0% - 1.7% = 1.3% [−0.4%, 3.0%], t-statistics = −1.11, p = 0.277), but these effects did not pass the statistical significance threshold after multiple testing correction) (Figure 3C; Table S3). In addition, we observed a reduced proportion of proliferative cells in Mac1 (13.1% - 35.0% = −21.9% [−21.1%, −22.7%], t-statistics = 2.79, p = 0.011) indicating that the cytokine mix may have an inhibitory effect on the proliferation of THP-1 macrophages. This is also in agreement with GSEA analysis conducted on a ranked transcriptome list (Table S1) from bulk RNAseq showing strong positive enrichment of DLAM gene sets and negative enrichment of proliferative gene sets (see Figure 2A), enrichment of Proliferative module scores (Figure 4C) and strong hypergeometric overlap between Mac1 cluster marker genes and Proliferative gene sets (Figure 4A).

To investigate the effects of cytokine mix treatment on gene expression, we performed differential expression analysis on pseudo-bulked cells using *edgeR* ^59^. The up-regulated DEGs from scRNA-seq showed strong concordance with the bulk transcriptomic data as evidenced by the rank hypergeometric overlap method (Figure S4B). The top up-regulated genes include *TGM2*, *SERPINB4* and *CCL5*, while top down-regulated genes include *IPCEF1* and *CD14*. The up-regulated genes show increased expression in two major DLAM-like clusters (Mac0, Mac4), whereas the top down-regulated genes have decreased expression in different DLAM-like clusters (Mac3, Mac7) (Figure 3D), suggesting that these THP-1 macrophage subtypes may be driving the transcriptional response to the cytokine mix exposure that we observe. Together, these findings indicate that cytokine mix-treated THP-1 macrophages can exhibit transcriptional profiles and states similar to those found in human AD brains and iMGLs treated with phagocytic, CNS-relevant substrates.

### Cytokine mix-treated THP-1 macrophages show a reduction in migration, phagocytosis, and lysosomal proteolysis

Phagocytic clearance of cellular debris rich in cholesterol and other lipids, such as dystrophic neurites, synapses, apoptotic cells, and myelin fragments (a.k.a. efferocytosis) is one of the biological processes that AD risk variants may affect to modulate disease susceptibility ^10–12^. This four-step biological process involves migration/recognition (“find-me”), engulfment (“eat-me”), degradation (“digest-me”) and adaptation/storage/elimination (“poop-me”) of such debris and the cholesterol/lipids derived from its digestion. (Figure 5A). In addition, global transcriptomic analysis showed that some aspects of efferocytic clearance may be increased including the degradation and adaptation processes (“Oxidative phosphorylation”, “Lysosome”, and increased DLAM signatures) and other aspects may be suppressed (“FC Gamma Receptor Phagocytosis”). Therefore, we functionally characterized the performance of THP-1 macrophages stimulated with the cytokine mix in each of the efferocytosis steps. First, we used a Scratch Wound Assay to evaluate macrophage motility and migration capacity. A scratch is performed in the center of the well and we imaged for 24 hours to quantify the cell density in the wound. We observed a reduction in the relative wound density in THP-1 macrophages stimulated with the cytokine mix compared to control THP-1 macrophages (Figure 5B) (Cytokine mix - control mean of differences (95% confidence interval) p=two-tailed paired t-test: −15.72 [−49.21 to 17.77] p=0.0181). This reduction in motility might be associated with a higher capacity for attachment that was observed when performing routine detachment of the macrophages by a trypsinization method in order to seed the cells for experimental procedures. Cytokine-induced macrophages also displayed significantly increased area as evidenced by increased confluence (Figure S5) In addition, we observed an increase in ICAM-1, an adhesion receptor, in cell media in THP-1 macrophages treated with the cytokine mix compared to control (Figure S6). Secondly, we investigated the phagocytic uptake ability of control and cytokine-stimulated macrophages using four different substrates. We used flow cytometry to quantify the uptake of fluorescent latex beads, pHrodo-labeled myelin, pHrodo-labeled zymosan, and pHrodo-labeled early apoptotic Jurkat cells (EAJ). We found uptake of all four phagocytic substrates was decreased as evidenced by lower phagocytic index (Figure 5B) Latex beads: −0.9171 [−1.135 to −0.6994] p=0.0009, Myelin: −1.654 [−2.429 to −0.8780] p=0.0065, Zymosan: −1.710 [−2.845 to - 0.5748] p=0.0173, EAJ: −3.745 [−5.899 to −1.591] p=0.0116).

**Figure 5.**
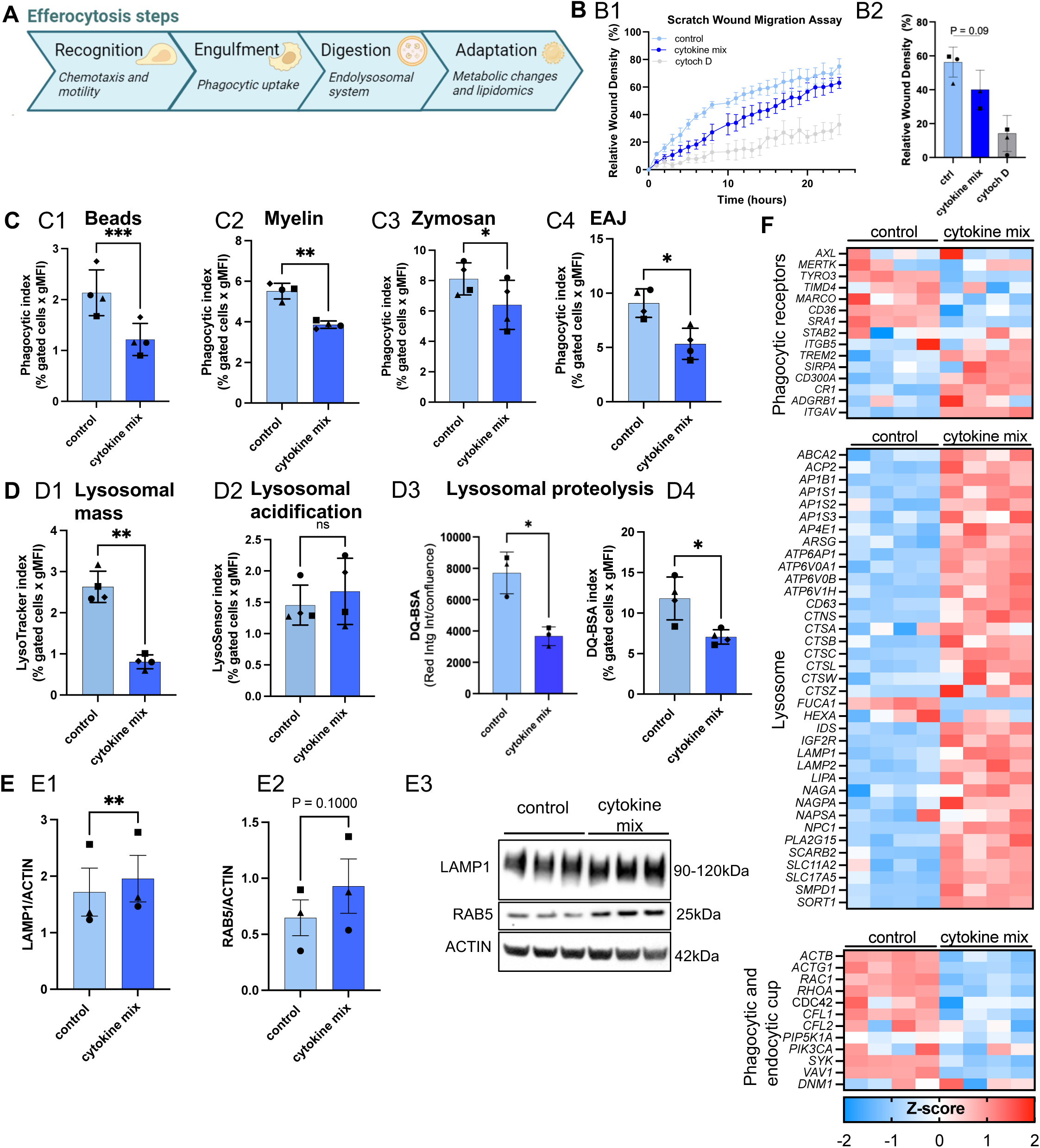
Cytokine mix treatment decreases phagocytosis and lysosomal processing in THP-1 macrophages. A) Diagram showing the steps in the efferocytosis process. B) Evaluation of the migration capacity through the scratch wound assay in THP1 macrophage control and treated with the cytokine mix. B1, relative wound density over time; B2, quantification of the area under the curve. Cytochalasin D (cytD) was used as a migration inhibitor. C) Quantification of phagocytic uptake of beads C1), myelin fragments C2), Zymosan C3), and early apoptotic cells (EAJ) C4). D) Evaluation of lysosomal activity in control and cytokine stimulated macrophages by LysoTracker (lysosomal mass, D1), LysoSensor (acidification, D2) and DQ-BSA (lysosomal proteolysis, D3-D4). Different dot shapes correspond to independent macrophage differentiations. N = 3-4 independent differentiation, each differentiation has 2-3 well replicates. Groups were tested with paired t.test (two-tailed) * p < 0.05, ** p < 0.01, *** p < 0.001.

To assess the degradative capacity of cytokine-treated macrophages, we evaluated lysosomal activity. We stained control and cytokine mix-stimulated THP-1 macrophages with markers for lysosomal mass (LysoTracker) and lysosomal acidification (LysoSensor). By using flow cytometry, we observed a significant reduction in the lysosomal mass as evidenced by lower lysosomal mass index: −1.823 [−2.617 to −1.028] p=0.0053, but no differences in lysosomal acidification represented as lysosomal acidification index: 0.2187 [−0.3352 to 0.7726] p=0.2979 in cytokine mix-stimulated THP-1 macrophages compared to control (Figure 5 D1, D2). Next, we exposed cells to DQ-BSA, a substrate used to evaluate lysosomal proteolysis. We found a reduction of DQ-BSA signal in THP-1 macrophages treated with the cytokine mix compared to control by time-lapse imaging as evidenced by lower Integrated Intensity normalized to Confluence: −4040 [−7664 to −415] p=0.0408 (Figure 5 D3). We confirmed decreased DQ-BSA signal 24h post DQ-BSA exposure in cytokine-treated macrophages by flow cytometry quantifying DQ-BSA index: −4.743 [−7.580 to −1.906] p=0.0130 (Figure 4 D4). This observation was unexpected because transcriptomic profiling of cytokine-treated macrophages showed up-regulation of genes involved in “Lysosomal” pathway (Figure 1C) such as *LAMP1, EEA1, RAB5* (Table S1), and cathepsins such as *CTSB, CTSD* (Figure 2B, Table S1) suggesting that lysosomal proteolytic capacity might be induced in cytokine-treated macrophages. Using western blot we confirmed that the LAMP1 and RAB5 proteins were up-regulated in cytokine-treated macrophages as evidenced by increased protein band signal normalized to Actin (LAMP1: 0.2383 [0.1708 to 0.3057], p=0.0043; RAB5: 0.2318 [−0.1334 to 0.6970], p=0.100) (Figure 5 E1, E2). To understand the discrepancies between functional response and transcriptomic changes in gene expression involved in lysosomal proteolysis and phagocytic uptake, we carefully examined the expression of genes involved in each step of phagocytic and endocytic clearance. We found that some phagocytic receptors were up-regulated such as (*TREM2* logFC = 0.52, adj.p = 0.0098; *CD300A* logFC = 1.36, adj.p = 9.77e-5; *ITGAV* logFC = 0.75, adj.p = 0.0003), while others were down-regulated (*TYRO3* logFC = −2.04, adj.p = 6.29e-06; *SRA1* logFC = −0.70, adj.p = 0.0001) in THP-1 macrophages exposed to cytokine mix (Figure 5F). We have also observed an increased expression of lysosomal genes such as (*ATP6AP1* logFC = 0.47, adj.p = 5.22e-07; *ACP2* logFC = 0.62, adj.p = 1.93e-06*; CTSB* logFC = 0.80, adj.p = 0.011; *SORT1* logFC = 0.95, adj.p = 1.06e-06) in THP-1 macrophages treated with cytokine mix (Figure 5F). Finally, we examined the expression of genes involved in formation of phagocytic and endocytic cups. This is an important step for both phagocytosis and endocytosis through which lipid-rich substrates and extracellular proteins such as BSA are internalized. We found that actin-related genes (*ACTB* and *ACTG1*) involved in regulation of actin polymerization as well as Rho family GTPases such as *RAC1* and *RHOA* that are essential for phagocytic/endocytic cup formation, are significantly down-regulated in cytokine-treated THP-1 macrophages which may explain reduced internalization of DQ-BSA and phagocytic substrates (*ACTB*, logFC = −0.232 adj.p = 0.0069; *ACTG1*, logFC = −0.239 adj.p = 0.0035; *RAC1*, logFC = −0.325 adj.p = 4.24e05; *RHOA*, logFC = −0.258 adj.p = 0.0036)(Table S1) (Figure 5F).

### Cytokine mix-treated THP-1 macrophages enhance oxidative and glycolytic metabolism

The uptake and digestion of whole cells or cellular fragments requires adaptation to the engulfed material, which includes transcriptional changes, lipid storage and efflux, and metabolic adaptation (Romero-Molina et al., 2023). Metabolic changes have been shown to be both a modulator and a consequence of the immune response in macrophages (Jha et al., 2015). We used the Seahorse technology to study glycolytic and mitochondrial bioenergetics. Upon stimulation with the cytokine mix, THP-1 macrophages increased basal: 128.7 [35.52 to 221.9] p=0.0085 and maximal respiration capacity: 251.9 [−10.64 to 514.4] p=0.0107 (Figure 6A). In parallel, our glycolytic stress test showed a pronounced increase in basal glycolysis: 90.83 [49.05 to 132.6] p=0.0002 and maximal glycolytic capacity: 53.18 [15.7 to 90.66] p=0.0079 (Figure 6B). These results demonstrate an enhancement of both oxidative and glycolytic metabolism in response to the cytokine mix treatment and are also in agreement with transcriptomic findings showing “Oxidative phosphorylation” and “Glycolysis Gluconeogenesis” as top positively enriched pathways (Figure 1C).

**Figure 6.**
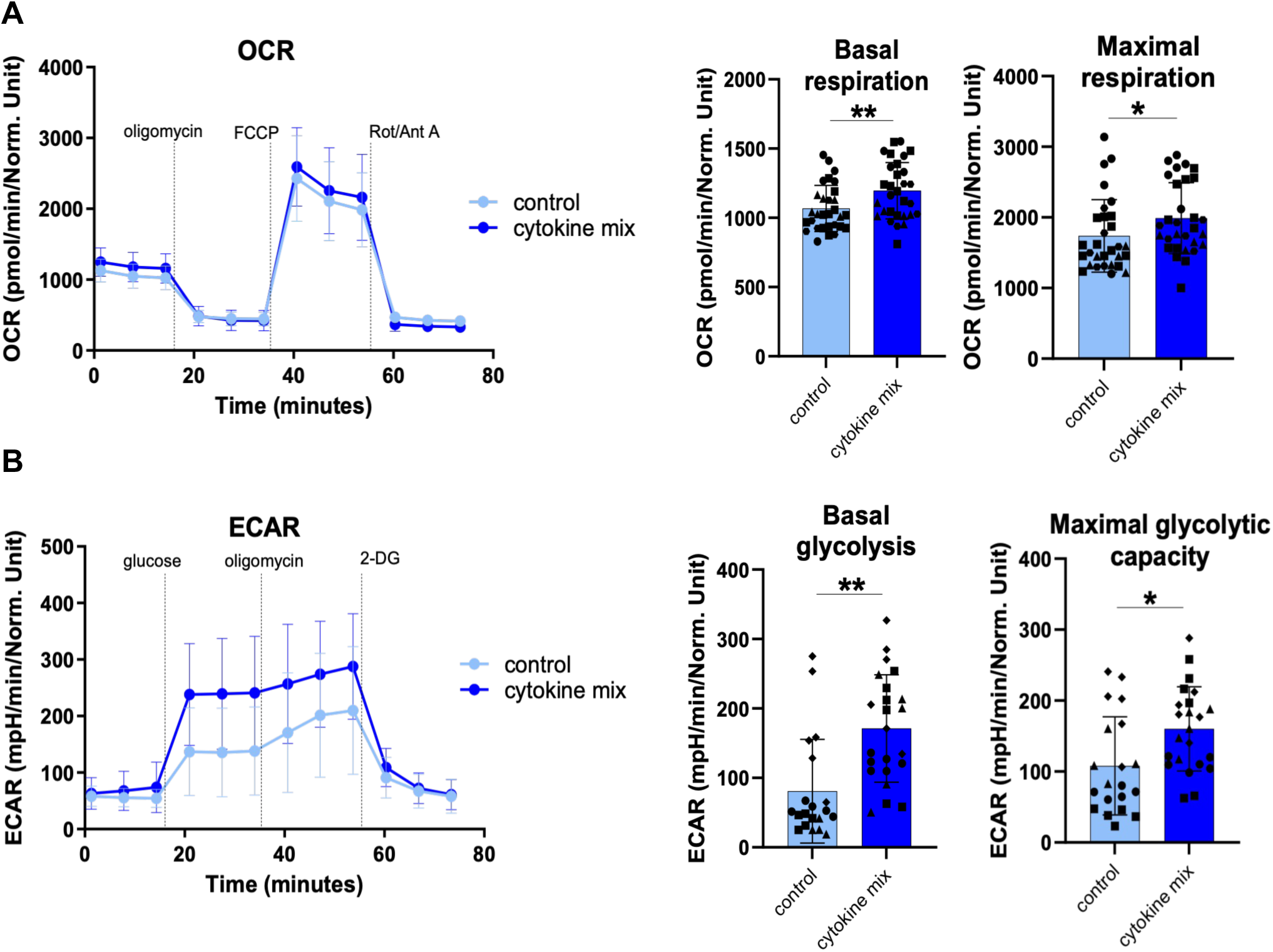
Cytokine mix treatment increases glycolytic and mitochondrial metabolism in THP-1 macrophages. A) Seahorse Mitostress Test: A1, Test profile of oxygen consumption over time. Quantification of basal (A2) and maximal (A3) respiration capacity. B) Seahorse Glycolysis Stress Test: B1, Test profile of extracellular acidification rate over time. Quantification of basal glycolysis (B2) and maximal glycolytic capacity (B3). Different dot shapes correspond to independent macrophage differentiations. N = 4 independent differentiation, each differentiation has 4-6 well replicates. Groups were tested with paired t.test (two-tailed) * p < 0.05, ** p < 0.01.

### Cytokine mix-treated THP-1 macrophages show significant changes in lipid metabolism

Lipid efflux and storage are essential features in the adaptation step of the efferocytosis process. Moreover, the accumulation of lipids in myeloid cells has been a subject of interest due to its potential role in the pathogenesis of AD and related neurodegenerative conditions ^60–63^.

Bulk RNA-seq analysis comparing THP-1 macrophages treated with the cytokine mix vs untreated controls showed statistically significant changes in the expression of lipid metabolism related genes (Table S1). There was an up-regulation of genes coding for proteins directly involved in cholesterol efflux and transport (*APOE* logFC = 1.74, adj.p = 0.003, *APOC1* logFC=3.03, adj.p=2.92e-06), hydrolysis of cholesterol esters in lysosomes (*LIPA*, logFC=2.12, adj.p=1.39e-08) or egress of cholesterol from lysosomes (*NPC2* logFC=0.40 adj.p=0.019) and cholesterol ester hydrolysis in lipid droplets (*NCEH1* logFC=0.96, adj.p=3.60e-05). In contrast, we observed a down-regulation of *SOAT1* (logFC=-0.33, adj.p=1.97e-04) involved in cholesterol ester synthesis and storage in lipid droplets.

To test whether these transcriptomic changes were associated with changes in lipid metabolism, we performed unbiased lipidomics in control and cytokine mix-treated THP-1 macrophages (Figure 7A). We observed a statistically-significant decrease in mono/di/tri-acylglyceride (MG, DG, TG) species after stimulation with the cytokine mix, which could be related to the increased energy expenditure observed in Seahorse experiments (Figure 7A). In addition, we observed a reduction in globotriaosylceramide (GB3), a glycosphingolipid related to lipid storage. On the other hand, we found a significant increase in lysophosphatidylethanolamines (LPE), phosphatidylethanolamines (PE), phosphatidic Acid (PA), and phosphatidylserine (PS), which might suggest membrane remodeling, such as increase in fluidity, or alterations in cell signaling pathways.

**Figure 7.**
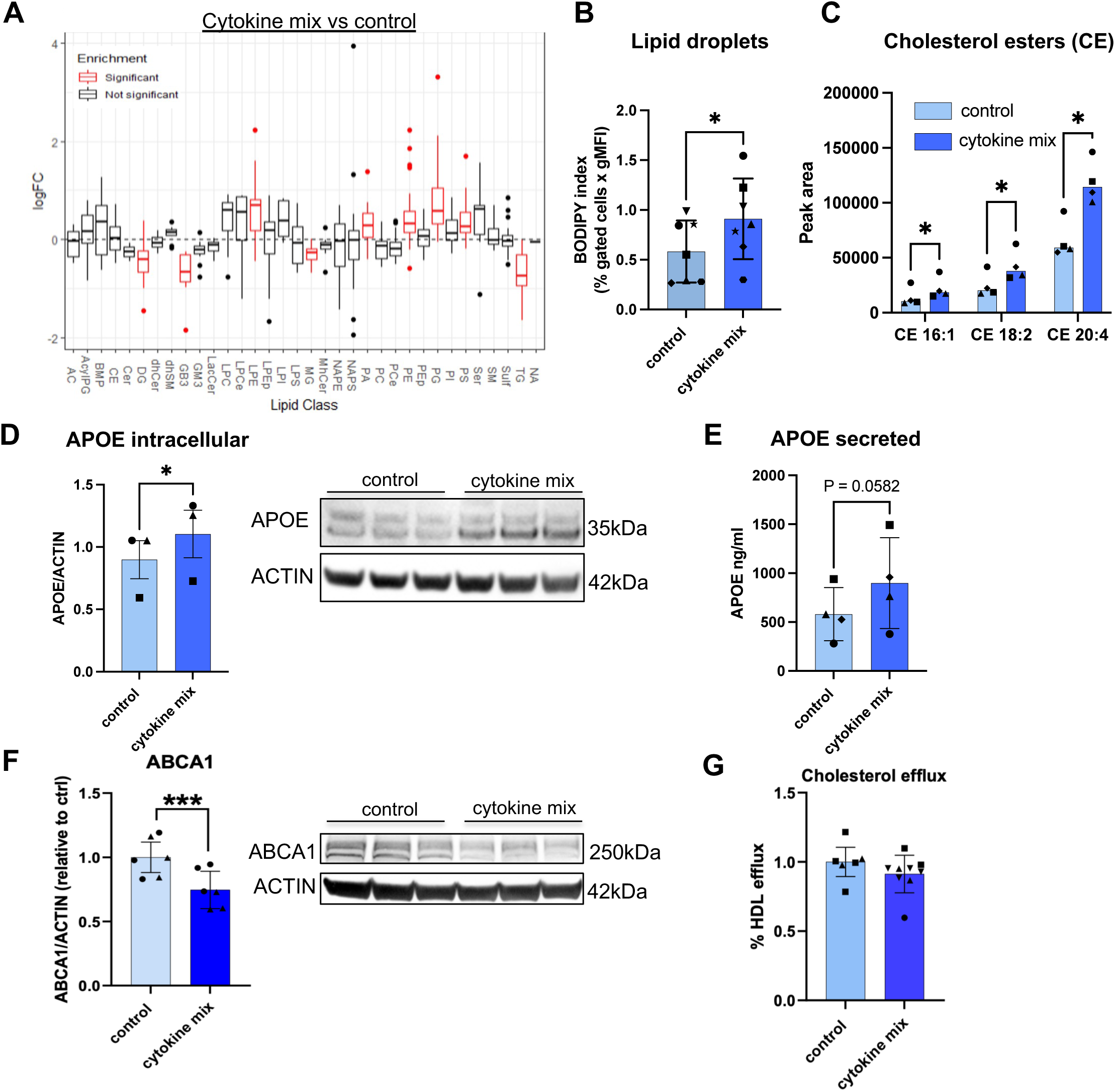
Cytokine mix treatment induces changes in lipidomic profiles and lipid metabolism in THP-1 macrophages. A) Lipidomics results from control and cytokine mix stimulated THP-1 macrophages (N=4). B) Quantification of lipid droplet content (BODIPY staining) by flow cytometry in THP-1 control and stimulated macrophages. C) Abundance of selected cholesterol ester species (raw data in Table S4). D) Quantification of intracellular APOE normalized to Actin in THP-1 macrophages treated with cytokines. Quantification (left), representative images (right). E) Quantification of secreted APOE in THP-1 macrophages. F) Quantification of ABCA1, normalized to Actin, measured by western blot in control and cytokine-treated THP-1 macrophages. Quantification (left), representative image (right) G) Quantification of cholesterol efflux, in THP-1 control and stimulated macrophages into HDL acceptor N = 3-4 independent differentiation, each differentiation has 3 well replicates. Different dot shapes correspond to independent macrophage differentiations. Groups were tested by paired t.test (two-tailed) * p < 0.05, ** p <0.01.

In addition, we used BODIPY to stain neutral lipids. By using flow cytometry, we observed increased BODIPY fluorescent signal in THP-1 macrophages treated with the cytokine mix as evidenced by increased BODIPY index: 0.32 [0.02 to 0.63] p=0.038 (Figure 7B). As mentioned above, the level of TGs (one of the lipid species that can be stored in lipid droplets) was decreased; however three cholesterol ester (CE) species were increased including CE 20:4 (logFC = 0.71, adj.p = 0.009), CE 18:2 (logFC = 0.66, adj.p = 0.01), and CE 16:1 (logFC = 0.56, adj.p = 0.034) (Figure 7C and Table S4). We then examined proteins involved in reverse cholesterol transport such as cholesterol acceptor APOE and cholesterol transporter ABCA1. Transcriptomic levels of *APOE* were markedly increased in cytokine mix-treated THP-1 macrophages compared to control (Table S1). We confirmed that cytokine mix-treated THP-1 macrophages express higher levels of APOE protein: 0.2060 [0.02506 to 0.3869], p=0.0392 (Figure 7D) and display increased levels of secreted APOE measured by ELISA (ng/ml): 317.3 [−20.55 to 655.1] p=0.0582 (Figure 7E). Surprisingly, we observed a decreased level of ABCA1 protein: −0.254 [−0.360 to −0.148], p=0.0003 (Figure 7F) as well as no changes in cholesterol efflux −0.111 [−0.266 to 0.044], p=0.137 (Figure 7G) suggesting that cholesterol efflux is not affected.

### Cytokine mix-treated THP-1 macrophages showed minor alterations in secreted cytokine profile

To test if the cytokine mix exposure is associated with a decrease in proinflammatory cytokines and induction of anti-inflammatory cytokines, we measured secreted cytokines in the conditioned media of THP-1 macrophages treated with cytokine mix and compared the levels of cytokines in the conditioned media of control cells. Using proteome profiler dot blot we found increased levels of secreted CCL2, CXCL1, ICAM1 and decreased levels of IL-1ra (Figure S6). *CCL2* is up-regulated in xMGL DAM (GS8) ^43,64^ and in human foamy macrophages (GS24) ^35^. Hence, elevated level of CCL2 expressed as relative density normalized to density of reference spot (0.518 [0.451 to 0.569], p<0.001), also markedly increased at the transcript level (Figure 2B), most likely indicates a higher DLAM-like population in cytokine mix-treated THP-1 macrophages. Consistently, the highest level of CCL2 expression was observed in the Mac0 cluster that shows the biggest enrichment with DLAM genesets (Figure S3A). Another cytokine induced by cytokine-mix treatment is CXCL1 as evidenced by increased relative density normalized to density of reference spot: 0.109 [0.046 to 0.173], p=0.012 which is a potent neutrophil chemoattractant ^65,66^.

## Discussion

Recent studies have focused on optimization of *in vitro* systems that recapitulate disease/lipid-associated microglia/macrophages. These models use iPSC-derived microglia, CNS-relevant substrates, or genetic manipulations of DLAM master transcription factors ^19,20,67^. Although they can model human microglia under controlled conditions and offer insights into microglial biology, they also have several caveats. For example, generation of iPSC-derived microglia is expensive and time consuming (often > 1 month). In addition, the use of different differentiation protocols and iPSC lines by different labs to generate iMGLs is a source of large variability ^68^. Yet, the efficiency to induce a DLAM-like state across these different methods and cells has never been fully investigated. In addition, to induce a DLAM state *in vitro* in HPC-derived iMGLs ^32^, one would need a combination of phagocytic substrates that have to be prepared in advance such as apoptotic cells, or myelin fragments and synaptosomes isolated from human or mouse brains^19^. Here, we propose use of an easily accessible and well-characterized human immortalized monocytic leukemia cell line (THP-1) that can be differentiated into macrophages and then stimulated with a cocktail of cytokines including maturation, survival and anti-inflammatory cytokines to induce a DLAM-like state *in vitro*.

Using bulk RNA-seq we found that THP-1 macrophages closely resemble iMGLs and *ex vivo* microglia making this cellular system relevant to investigate brain-resident macrophages. We demonstrated that a DLAM-like transcriptional response can be induced *in vitro* in THP-1 macrophages with a mixture of four cytokines as evidenced by activation of similar transcriptional programs to those found in AD human brains ^36,44–47^ and iPSC-derived microglia exposed to CNS-derived lipid-rich substrates ^19^. Consistently, single cell RNA-seq analysis of control and cytokine mix-treated THP-1 macrophages identified 9 unique clusters enriched for DLAM, proliferative and antigen-presenting gene signatures from published studies ^15,16,19,45,47,48^ and related biological functions. Interestingly, projection of DLAM clusters identified in this study to two DAM clusters induced by phagocytic substrates in iMGLs showed that our Mac DLAM clusters closely resemble Cluster 8 DAM identified by Dolan et al. that appear upon stimulation with either Aβ fibrils or apoptotic cells ^19^. Of note, both module score analysis and cluster projection using the *scmap-cluster* package, clearly showed that DLAM exists at baseline in control macrophages. This might be induced by a local turnover of macrophages when dying (apoptotic or necrotic) cells need to be phagocytosed and cleared. However our study showed that the proportion of clusters (Figure 3D, Figure S3D) can be shifted toward more DLAM, less proliferative by using a combination of cytokines. Importantly, we identified that several DLAM clusters match the transcriptional signature of microglia found in AD brains such as lipid processing microglia or GPNMB^+^ DAM clusters from AD biopsies^45^. For example Mac6 showed strong enrichment of lipid processing cluster marker genes (MG4) ^47^ suggesting that we can model some of the *in vivo* microglial states in a simple *in vitro* system.

We also demonstrated that IL4 is the main driver of increased DLAM transcriptional responses and reduction of proliferative responses in our cytokine mix, while 3F cytokines have no impact on any of these transcriptional programs. Similarly, IL4 induced the expression of AD risk genes in the same direction as in the cytokine mix with the exception of *MS4A4A*, *MS4A6A*, and *PILRA* that were induced in the opposite direction in cytokine mix and 3F suggesting that MCSF, IL34, and/or TGFβ control the expression of these three AD risk genes. Altogether, we identified a mix of four cytokines (IL4, MCSF, TGFβ, IL34) that may be a useful tool to manipulate macrophage states *in vitro* toward a more AD-relevant state; which so far can be achieved by more expensive and challenging methods such as exposure of iMGLs to synaptosomes, myelin fragments, and apoptotic neurons ^19^, reduction of transcription factors (BHLHE40/41) that repress DLAM responses ^20^, and overexpression of a positive DLAM regulator, MITF^19^.

Another goal of this work was to functionally characterize DLAM-like cells *in vitro* in order to link the biological pathways associated with DLAM states defined by the transcriptomic analyses with experimental measures of their activity. The main DLAM-enriched processes are phagocytosis, lipid and lysosomal clearance and modulation of immune response ^15,16^. We have found that cytokine-stimulated macrophages showed decreased uptake of beads and decreased phagocytosis of cholesterol and lipid rich cellular debris, including early apoptotic cells, myelin and zymosan. The consequences of the polarization towards DLAM state on phagocytosis are not clear yet. On one hand, a recent study showed that overexpression of DLAM-regulating transcription factor, MITF, leads to increased phagocytic uptake of myelin ^19^. On the other hand, reduction of *SPI1* (encoding PU.1), also associated with delayed AD onset ^4^ led to decreased phagocytosis of myelin and zymosan ^69^. Although an enhanced phagocytic capacity may be beneficial for increased debris clearance, a reduced uptake may be advantageous in AD, considering how microglial uptake of cholesterol/lipid-rich cellular debris during neurodegeneration may result in microglial lipid overload and dysfunction, leading to cell senescence and death or polarization toward a neurotoxic, proinflammatory or otherwise detrimental state. Increased phagocytic uptake may also be a symptom of dysfunctional lipid trafficking to lysosomes in microglial cells as was previously described in Niemann-Pick type C neurodegenerative disease ^70^. In addition, it may slow down the removal of synapses observed in neurodegenerative diseases. Finally, in our system, decreased phagocytosis may rather be the effect of reduced actin polymerization and phagocytic cup formation than decreased efficiency of the endolysosomal system.

The final step of efferocytosis involves metabolic and other molecular and cellular adaptations, as well as storage and elimination of cholesterol and other lipids and molecules derived from the digestion of engulfed material. This step is particularly important to handle the excess cholesterol derived from the degenerating brain, the most cholesterol-rich organ in the body. Cytokine mix-treated THP-1 macrophages showed an increased level of lipid droplets, APOE expression and secretion, but decreased expression of ABCA1 and no change in cholesterol efflux. Although we observed induced expression of DLAM genes involved in endolysosomal processing and cholesterol clearance, we also found an increased level of two DLAM inhibitors, *BHLHE40* and *BHLHE41* (Table S1), that compete with LXR and MIT/TFE family members to decrease DLAM responses. We have previously shown that loss of BHLHE40/41 is associated with increased lysosomal and lipid clearance ^20^. Even though the majority of DLAM responses are induced at the transcript level, functional responses in cytokine-treated THP-1 macrophages may be partially inhibited due to BHLHE40 and BHLHE41 that inhibit positive transcriptional regulators of the DLAM response such as LXR, PPAR, and MITF/TFEB family TFs. In addition, BHLHE40 has been nominated by us as a regulator of mouse DLAM ^20^, and its induction has been shown to be a part of mouse DAM response ^17,42^.

Macrophages, including microglia, demonstrate remarkable plasticity, adapting their metabolism to meet the specific demands of different microenvironments and immune challenges. In the pro-inflammatory activation state (M1), macrophages predominantly rely on glycolysis to rapidly generate energy and produce pro-inflammatory cytokines, while in the anti-inflammatory activation state (M2), macrophages exhibit a shift towards oxidative phosphorylation, promoting tissue repair ^71^. However, it has recently been shown that a pro-inflammatory treatment (LPS) for 24 hours increases both glycolytic and oxidative metabolism in HMC3 microglia ^72^. Similarly, our results showed an enhancement of both oxidative and glycolytic metabolism, which is in accordance with the mix of M1/M2 transcriptional profiles found in our cytokine-treated THP1 macrophages ^30^. The fact that the increase in basal glycolysis had a higher effect size than the increase in basal mitochondrial respiration may be explained by the high glucose availability present in the media *in vitro,* or be a feature of the DLAM transcriptional profile, resembling the Warburg effect described in T cells (aerobic glycolysis without cell proliferation) ^73^. APOE4 primary mouse microglia, which display an up-regulation of DAM genes compared to APOE3 microglia, show higher rates of basal glycolysis and lower maximal respiration than APOE3 microglia ^74^. However, *in vivo* data in AD mouse models showed a major up-regulation of oxidative phosphorylation in active (Clec7a^+^) microglia ^75^. The transmembrane receptor TREM2 is essential for polarization towards a DLAM state ^15^, as well as for microglial metabolic fitness ^76^, supporting the link between the transcriptomic and metabolic changes underlying the immune response. Our results showed that the transcriptomic changes that occur when macrophages are polarized towards a DLAM phenotype enhance both their glycolytic and oxidative metabolism to increase energy production.

Triacylgliceride (TG) synthesis has been proposed to enhance macrophage inflammation ^77^. *In vitro,* mouse pro-inflammatory macrophages (LPS+IFNγ stimulation) increase TGs, in contrast to mouse macrophages treated with an anti-inflammatory stimulus (IL4), which showed a reduction in some TGs compared to control macrophages ^77^. In concordance, our results showed a decrease in TGs in human macrophages treated with the cytokine mix, which contains IL4. On the other hand, upon chronic demyelination (12 weeks cuprizone treatment), *Trem2-KO* mouse microglia showed an increase in triacylglycerol (TG) and ganglioside (GB3) compared to WT mouse microglia, suggesting that TREM2 might modulate lipid metabolism ^24^. THP-1 macrophages treated with cytokine mix showed an increase in *TREM2* expression coupled with a decrease in TG and GB3, compared to control macrophages, reinforcing the idea that TREM2 might play a role in immunometabolism ^24,76^.

We also observed that the polarization towards a disease-associated state upon cytokine mix treatment induced an increase in lipid droplets (LD) in THP-1 macrophages. In accordance, Claes et al., showed lipid droplet accumulation in human DAM xenotransplanted microglia in the brain of chimeric AD mice ^64^. More interestingly, xenotransplanted TREM2-R47H mutant human microglia exhibited a reduction in the accumulation of lipid droplets *in vivo* ^64^. Prakash et al. showed that the lipid droplet load in microglia was positively correlated with their proximity to amyloid plaques in the 5xFAD mouse model and the human brain ^78^. In addition, LD-laden microglia showed a deficit in Aβ phagocytosis ^78^, which may explain the decreased phagocytic capacity that we observed in cytokine mix-treated THP1 macrophages. Finally, we found an increased level of CXCL1 that leads to increased accumulation of oxLDL and increases the rate of conversion to foam macrophages ^79^ which may also explain the increased level of lipid droplets in cytokine mix treated THP-1 macrophages. On the other hand, Haney et al recently described LD-associated microglia in APOE4 carriers ^63^. These toxic microglia, previously found by the same laboratory in aged mice ^62^ are transcriptionally distinct from DLAM ^20^ and accumulate TGs rather than CEs in LD. Isolated microglia from APP/PS1:APOE4 mice displayed an increase in both TGs and CEs compared to microglia from APP/PS1:APOE3 mice^80^. Given a decreased level of several TGs species, LD in cytokine-treated THP-1 macrophages may be associated with non-inflammatory and protective rather than the detrimental phenotype.

In conclusion, we have described here a convenient tool to polarize THP-1 macrophages toward a DLAM state *in vitro*, using a cocktail of anti-inflammatory and macrophage maturation cytokines. We performed bulk and single cell transcriptional profiling of cytokine mix-treated THP-1 macrophages and found robust activation of DLAM transcriptional responses as well as manipulation of the expression of several AD risk genes. The use of a human immortalized cell line may constitute an advantage in terms of time, cost and replicability compared to the use of induced pluripotent stem cells. Finally, since our model is more easily manipulated in functional genomic studies, we performed a broad characterization that provides insight into the function of macrophages with increased expression of DLAM genes showing lower phagocytosis and lysosomal proteolysis, higher oxidative and glycolytic metabolism and lipid modifications. Future studies should confirm these phenotypes *in vivo*, in the presence of AD and other pathologies (e.g., age-related demyelination ^81^) and coupled with interactions with other cell types.

## Data and code availability

All aligned read counts and fastQ files for cytokine-treated and control THP-1 macrophages from bulk and scRNAseq studies have been deposited to the Gene Expression Omnibus and are available under the following accession numbers GSE273482 (bulk RNAseq), GSE273912 (scRNAseq). Additionally, TPM, DGEA and GSEA are listed in Table S1. Cluster marker genes and proportion between clusters related to scRNAseq analysis are listed in Table S3. Lipidomics raw data are available in Table S4 accompanying this manuscript.

## Acknowledgments

This work was funded by grants from the NIH: RF1AG054011 (A.M.G.), U01AG058635 (A.M.G., E.M.), R56AG081417 (E.M., A.M.G.), The JPB Foundation (A.M.G.), and The Belfer Neurodegeneration Consortium (A.M.G., E.M.). C.R-M. acknowledges support from the Alfonso Martin Escudero Postdoctoral Fellowship and the Alzheimer’s Association (24AARF-1191936). A.P-D. received funds from BrightFocus Foundation A2024025S and Training Program in Stem Cell Biology fellowship from the New York State Department of Health (NYSTEM-C32561GG). This work was supported in part through the computational and data resources and staff expertise provided by Scientific Computing and Data at the Icahn School of Medicine at Mount Sinai and supported by the Clinical and Translational Science Awards (CTSA) grant UL1TR004419 from the National Center for Advancing Translational Sciences.

## Author contributions

A.P-D, C.R-M, E.M, A.M.G - conceptualization;, A.P-D, C.R-M, T.P, E.M, A.M.G - manuscript draft and editing; A.P-D, C.R-M, W.Y.S - data acquisition and analysis, T.P., A.P-D - single cell data analysis, Y.L. - bulk RNAseq processing.

## Materials and Methods

### THP-1 differentiation and cytokine treatment

THP-1 monocytes were cultured in RPMI medium supplemented with 10% FBS, 1x Penicillin Streptomycin (1,000U/ml Penicillin 1,000μg/ml Streptomycin) and 10mM HEPES. 25ng/ml of phorbol 12-myristate 13-acetate (PMA) was used to differentiate THP-1 monocytes to macrophages. After 3 days, PMA was removed and replaced with PMA-free RPMI media (as described above). THP-1 macrophages were treated with 20ng/ml IL-4, 25ng/ml MCSF, 50ng/ml TGFβ and 100ng/ml IL34 for 2-3 days and cells were collected, or functional assays were performed.

### Cholesterol efflux assay

Cholesterol efflux was performed using a Cholesterol Efflux Fluorometric Assay kit (Biovision, K582-100) following the manufacturer’s instructions. For this assay, cells were seeded in a 96-well plate at 50,000 cells/well. Cells were labeled with a Labeling Reagent for 1h at 37°C followed by loading cells with an Equilibration Buffer. After overnight incubation, media containing Equilibration Buffer was aspirated and replaced with media containing cholesterol acceptor human HDL (40 µg/ml) for 5h at 37°C. At the end of the incubation, supernatants were transferred to flat bottom clear 96-well white polystyrene microplates (Greiner Bio-one, 655095). Adherent cell monolayers were lysed with Cell Lysis Buffer and incubated for 30 min at RT with gentle agitation followed by pipetting to disintegrate cells. Cell lysates were transferred into flat bottom clear 96-well white polystyrene microplates. Fluorescence intensity (Ex/Em=485/523nm) of supernatants and cell lysates was measured using a Varioskan LUX multimode microplate reader (Thermo Fisher Scientific, VL0000D0). Percentage of cholesterol efflux was quantified: % cholesterol efflux = Fluorescence intensity of supernatant / fluorescence intensity of supernatant plus fluorescence intensity of cell lysate x 100.

### Lipidomic analysis

THP1 macrophages were seeded in a 6-well-plate (1,000,000 cells/well) and differentiated to macrophages. Control and cytokine-stimulated macrophages were collected after 2 days of treatment and cell pellets were shipped to Columbia University Biomarker Core (NY, US), where a standard lipid panel was performed. We performed 4 independent macrophage differentiations, and each sample represented cells pooled from 3 wells (a total of 3,000,000 cells per sample).

Lipidomics profiling was performed using Ultra Performance Liquid Chromatography-Tandem Mass Spectrometry (UPLC-MSMS) (Agudelco et al 2020, Area-Gomez et al 2021) in collaboration with the Core at Columbia University. Lipid extracts were prepared from cell lysates spiked with appropriate internal standards using a modified Bligh and Dyer method and analyzed on a platform comprising Agilent 1260 Infinity HPLC integrated to Agilent 6490A QQQ mass spectrometer controlled by Masshunter v 7.0 (Agilent Tecmethod and Santa Clara, CA). Glycerophospholipids and sphingolipids were separated with normal-phase HPLC as described before (Chan et al, 2012), with a few modifications. An Agilent Zorbax Rx-Sil column (2.1 x 100 mm, 1.8 µm) maintained at 25°C was used under the following conditions: mobile phase A (chloroform: methanol: ammonium hydroxide, 89.9:10:0.1, v/v) and mobile phase B (chloroform: methanol: water: ammonium hydroxide, 55:39:5.9:0.1, v/v); 95% A for 2 min, decreased linearly to 30% A over 18 min and further decreased to 25% A over 3 min, before returning to 95% over 2 min and held for 6 min. Separation of sterols and glycerolipids was carried out on a reverse phase Agilent Zorbax Eclipse XDB-C18 column (4.6 x 100 mm, 3.5um) using an isocratic mobile phase, chloroform, methanol, 0.1 M ammonium acetate (25:25:1) at a flow rate of 300 μl/min. Quantification of lipid species was accomplished using multiple reaction monitoring (MRM) transitions (Chan et al, 2012; Hsu et al, 2004; Guan et al, 2007) under both positive and negative ionization modes in conjunction with referencing of appropriate internal standards: PA 14:0/14:0, PC 14:0/14:0, PE 14:0/14:0, PG 15:0/15:0, PI 17:0/20:4, PS 14:0/14:0, BMP 14:0/14:0, APG 14:0/14:0, LPC 17:0, LPE 14:0, LPI 13:0, Cer d18:1/17:0, SM d18:1/12:0, dhSM d18:0/12:0, GalCer d18:1/12:0, GluCer d18:1/12:0, LacCer d18:1/12:0, D7-cholesterol, CE 17:0, MG 17:0, 4ME 16:0 diether DG, D5-TG 16:0/18:0/16:0 (Avanti Polar Lipids, Alabaster, AL). Lipid levels for each sample were calculated by using peak area. LipidR package (version 2.16.0, https://bioconductor.org/packages/lipidr)^88^ in R studio software (R Core Team (2023). A Language and Environment for Statistical Computing. R Foundation for Statistical Computing, Vienna, Austria; https://www.R-project.org/) was used to graph and analyze the data.

### Lipid droplet assay

Lipid droplet (LD) quantification was performed using FACS. Cells were collected and stained with 3.7 µM BODIPY for 30 minutes at room temperature (RT), protected from light. For FACS, single-cell data were acquired using Attune flow cytometer (Thermo Fisher Scientific) and analyzed using FCS Express 7 (De Novo Software). Gates were set up based on fluorescence minus one (FMO) controls. BODIPY index was quantified to take into account both percentage of positive cells and changes in geometric mean fluorescent intensity (gMFI) (% of positive cells x gMFI)/10^5^.

### Lysosomal assays

THP-1 macrophages were incubated with 75 nM LysoTracker-Red (Thermo Fisher Scientific, L7525) for 20 min at 37°C followed by 1 µM LysoSensor-Green (Thermo Fisher Scientific, L7535) staining for 1 min at 37°C. To characterize hydrolytic capacity of lysosomes, cells were treated with 1µg/ml DQ Red BSA (Thermo Fisher Scientific) for 1h at 37°C. After collecting the cells, single-cell data were acquired using Attune flow cytometer (Thermo Fisher Scientific) and analyzed using FCS Express 7 (De Novo Software). Gates were set up based on fluorescence minus one (FMO) controls. To quantify lysosomal activity, LysoTracker index, Lysosomal index and DQ-BSA index were calculated analogously to how we quantify phagocytic index as a measure of phagocytic activity^4^. These metrics take into account both percentage of positive cells and changes in geometric mean fluorescent intensity for each marker (gMFI) (% of positive cells x gMFI)/10^5^. Additionally, DQ-BSA red fluorescent signal was quantified over time using the Incucyte S3 live imaging system. Cells were plated in 96-well plates (40,000 cells/well) and treated with 1 µg/ml of DQ-BSA. Images were acquired every hour over 5h at 37°C. Total integrated density was calculated as mean red fluorescence intensity multiplied by surface area of masked object [RCU x µm^2^] and normalized by cell confluence (phase channel).

### Migration assay

The Incucyte Scratch Wound Assay (Sartorius) is a real-time and automated method for studying cell migration and wound healing. THP1 monocytes (50,000 cells/well) were seeded in a 96-well tissue culture plate and differentiated to macrophages. A scratch/wound was generated in the cell monolayer using a 96-pin WoundMaker tool. Two washes were performed to eliminate cell debris. The plate was then placed into the Incucyte Live-Cell Imaging System, where images were captured every hour. The Incucyte software analyzed the images and enabled the quantification of wound closure over time. We performed 3-4 independent experiments with 3-4 technical replicates per experiment that were averaged within each experiment.

### Phagocytosis assays

Uptake of different bioparticles was measured by flow cytometry (FACS) in control or cytokine-stimulated THP-1 macrophages. The following bioparticles were used: human myelin fragments (10 μg/ml), early apoptotic Jurkat cells (EAJ) 1xEAJ/THP-1 macrophage, 5 μg/ml zymosan particles (Thermo Fisher Scientific, cat. P35364) or Carboxyl Fluorescent Polystyrene 1.0μm particles (CD Bioparticles cat. DCFG-L007) for 24h. Myelin fragments were isolated from human brain tissue (corpus callosum) as described previously^89^. Early Apoptotic Jurkat cells (EAJ) were prepared using 3-hour incubation with 1 μM staurosporine (Alfa Aesar, cat. J62837-M) as described previously^69^. Isolated myelin fragments and EAJ were labeled with pHrodo dye (Thermo Fisher Scientific, cat. P36600) (10μg/ml) in PBS for 30 min in the dark at room temperature followed by two washes in PBS. To inhibit phagocytosis cells were pre-treated with 2 μM Cytochalasin D (Sigma-Aldrich, cat. C8273-1MG) for 30 min and during incubation with bioparticles. After 24-hour incubation cells were collected with trypsin (Gibco, cat. 25200), washed twice, resuspended in 1.0% PBS/bovine serum albumin (BSA) buffer and analyzed on an Attune flow cytometer (Thermo Fisher Scientific). Cells were pre-stained with Live/Dead Fixable Violet Dead Cell Stain kit (Life Technologies, cat. L34955) to exclude dead cells and CD11b-FITC antibody (BioLegend, cat. 101206) for 30 min and then used for FACS. Data were analyzed using FCS Express 7 (De Novo Software). Live/Dead^Neg^CD11b-FITC^+^/pHrodo^+^ cells were used to quantify the phagocytic index: percentage of pHrodo^+^ cells in CD11b-FITC^+^ gated population x geometric mean pHrodo intensity / 10^6^; and represented as phagocytic activity as previously described^4^. Gates were set up based on fluorescence minus one (FMO) controls.

### Assessment of disease-associated surface markers by flow cytometry

Control or cytokine treated THP-1 macrophages were first washed with PBS then dissociated using Accutase for 10 min at 37°C using. Cells were spun at 400×g for 5 min and resuspended in 1.0% PBS/bovine serum albumin (BSA). Cells were pre-stained with the following dyes/antibodies: Live/Dead Fixable Violet Dead Cell Stain kit (Life Technologies, cat. L34955) to exclude dead cells, CD11b-FITC (BioLegend, cat. 101206) and one of the DAM markers CD11c-APC 1:100 (BioLegend, cat. 301614), CD63-APC 1:100 (BioLegend, cat. 353007), CXCR4-APC 1:100 (R&D Systems, cat. FAB172R), CD44-APC 1:100 (BioLegend, cat. 338805), CD274-APC 1:100 (BioLegend, cat. 329707). Only one DAM marker was used at a time, DAM-specific antibodies were not mixed at any time. Cells were washed twice in 1.0% PBS/BSA buffer and analyzed by Attune flow cytometer (Thermo Fisher Scientific). Gates were set up based on fluorescence minus one (FMO) controls. Data were analyzed using FCS Express 7 (De Novo Software).

### Seahorse experiments and mitochondrial assays

THP-1 monocytes were plated at 50,000 cells per well in a Seahorse 96 well-plate. Agilent Seahorse Mito Stress Test Kit (103015-100) was used to assess mitochondrial respiration. DMEM XF Base Media was supplemented with 1mM pyruvate, 2mM glutamine and 10mM glucose. Agilent Seahorse Glycolysis Stress Test Kit (103020-100) was used to address glycolytic capacity. DMEM XF Base Media was supplemented with 2mM glutamine. For both assays, cell media was replaced with 180ul of specific media and cells were incubated for 1h at 37°C in a non-CO2 incubator.

### Transcriptomic analysis

RNA from human THP-1 macrophages (MACs was extracted using the RNeasy Plus Mini kit (Qiagen, 74136) following manufacturer’s instructions. mRNA quantity was measured using Nanodrop 8000 (Thermo Fisher Scientific). RNA was submitted to Azenta (New Jersey, NJ, USA) for QC, library preparation, and next-generation sequencing. Samples passed quality control with Qubit and BioAnalyzer showing RIN > 9.0. Libraries were prepared using TruSeq RNA Sample Prep Kit v2 and paired-end sequenced using HiSeq2500 at a read length of 150bp to obtain 20-30M mapped fragments per sample. Sequenced reads were assessed for quality (FastQC v0.11.8), trimmed for adapter contamination (Cutadapt v2.6), and aligned to the human genome hg38 (STAR v2.5.3a). Differential gene expression analysis (DGEA) was performed using a linear mixed model implemented in DREAM (differential expression for repeated measures, variancePartition R package v1.23.1 and R v3.5.3^86^). Genes with FDR adj.p < 0.05 were considered differentially expressed (DEGs). To identify pathways enriched in THP-1 macrophages treated with the cytokines, we used Gene Set Enrichment Analysis (GSEA). Briefly, ranked lists were generated from differential gene expression analyses by ordering genes according to the signed Z statistic (Table S1). Ranked lists were analyzed using the “GSEA Preranked” module using default GSEA settings including 1000 permutations. Our preranked lists were tested for enrichment against genesets from the Molecular Signatures Database (MSigDB v7.5.1, Broad Institute, c5.bp.v2023.2.Hs.symbols.gmt, c5.mf.v2023.2.Hs.symbols.gmt, c5.cc.v2023.2.Hs.symbols.gmt, c2.cp.reactome.v2023.2.Hs.symbols.gmt, and c2.cp.kegg_legacy.v2023.2.Hs.symbols.gmt). DGEA and GSEA results are shown in Table S1. In addition, we tested for enrichment of DLAM gene sets selected from several publications. Gene sets and their sources are listed in Table S2. Signatures extracted from mouse datasets were lifted to human orthologs using the Orthology search tool in gProfiler^90^. Enrichment scores were normalized by geneset size to generate normalized enrichment scores (NES) according to the standard protocol^91^.

### Rank-rank hypergeometric overlap (RRHO)

Transcriptional signatures from bulk RNAseq (Table S1) and pseudo bulk scRNAseq (Table S3) were compared pairwise using the RRHO2 R package^87^. The recommended -log10(P-value) * sign(log2FC) metric was used to generate ranked lists of genes for each transcriptional signature. RRHO2 was then used to visualize both concordant and discordant gene expression changes across each pair of signatures as rank-rank hypergeometric overlap (RRHO) heatmaps. The color temperature of each pixel in an RRHO heatmap represents the negative log10-transformed hypergeometric overlap test P-value of subsections of the two ranked gene lists, adjusted for multiple testing using the Benjamini-Hochberg correction method. Heatmaps generated using RRHO2 have top-right (both decreasing) and bottom-left (both increasing) quadrants, representing concordant gene expression changes, while the top-left and bottom-right quadrants represent discordant gene expression changes. The default step size and p representation method (hyper) were used.

### Single-cell RNAseq library preparation

Control or cytokine treated THP-1 macrophages were first washed with PBS, then trypsinized for 10 min at 37°C (Gibco, cat. 25200). Trypsinization was halted by the addition of an equal volume of warm RPMI medium with 10mM HEPES and 10% FBS, cells were spun at 400×g for 5 min and resuspended in 1.0% PBS/bovine serum albumin (BSA). Cells were filtered through 40μm cell strainer to exclude cell clumps. Trypan blue was used to assess cell number and viability (typicallyY>Y90%) using an automated cell counter Countess (Thermo Fisher Scientific). Gel bead-in-emulsion (GEM) encapsulation and single cell indexing reactions were performed using a Chromium X™ Controller instrument (10× Genomics, Inc., Cat. No. 1000202). Single-cell 3’ RNA-seq libraries were prepared using the Chromium Next GEM Single Cell 3’ GEM, Library and Gel Bead Kit (version 3.1 chemistry, 10× Genomics, Inc., Cat. No. 1000121), according to the manufacturer’s instructions. Samples were processed in two independent batches. Libraries were sequenced on a NovaSeq 6000 System, with paired end 150Y×Y150 bp sequencing.

### Single cell RNAseq data analysis and integration

Raw base call files from the sequencer were demultiplexed into FASTQ files using the CellRanger workflow v7.1.0 (10x Genomics). FASTQ files were aligned to the human reference genome (GRCh38) followed by filtering, barcode counting and UMI counting to generate a feature-barcode matrix per sample. Quality control, normalization, clustering and marker gene identification were performed with Seurat v5.0.2^92^. Briefly, cells were removed if they were expressing fewer than 100 genes, less than 500 UMIs and greater than 95% of total UMIs, or if greater than 10% of reads mapped to the mitochondrial genome. Raw count data was normalized using SCTransform v0.4.1^93^, with regression for mitochondrial mapping percentage. Principal components analysis (PCA) was performed to determine the top most variable genes and generate PCs. Samples were integrated using harmony v1.2.0^83^ to account for batch differences. Clustering and data reduction were performed in Seurat using UMAP for visualization. The number of cells per batch were randomly downsampled to the smallest treatment condition to ensure that differences in proportions were not due to variable cell counts. Marker genes for each cluster were identified using Seurat FindMarkers. Cluster annotations were determined using a combination of approaches, (1) high expression of established marker genes from the literature; (2) enrichment of myeloid gene expression signatures from published datasets (Table S2) using hypergeometric overlap; (3) enrichment of biological pathways in the expressed genes to determine functional significance. We used the propeller method via the speckle package in R to identify statistically significant differences in cell proportions between treatment conditions for each cluster, with adjustment for experimental batch and using adj.p < 0.05 as a cutoff. For pseudobulk differential gene expression analysis (DGEA) between conditions, cells were aggregated to obtain average expression per gene per independent run (N=2 per condition) using Seurat AverageExpression function. Pseudobulked DGEA was performed using the edgeR^59^ package in R, with adjustment for the experimental batch to account for technical variability. For comparison of clusters identified in this study with iMGL clusters we used the scmap package (version 1.20.0)^85^. Briefly, Seurat objects were converted to SingleCellExperiment objects (SingleCellExperiment package, version 1.20.1). Projection of query (this study) and reference clusters^19^ was performed using scmap-cluster projection with a default feature selection (n_feature = 500). Enrichment of transcriptional programs was tested using the AddModuleScore function in Seurat. Transcriptional signatures are listed in Table S2.

### Western blotting

Cells were lysed in RIPA buffer (Thermo Scientific, 89900) supplemented with Protease/Phosphatase Inhibitor Cocktail (Cell Signaling, 5872) following manufacturer’s instructions. Protein concentration was measured using BCA kit (Thermo Fisher Scientific, 23225) and equal quantities were used to prepare samples for western blotting. Samples were resolved by electrophoresis with Bolt 4–12% Bis-Tris Plus Gels (Invitrogen) in Bolt MES SDS running buffer (Invitrogen, B0002) and transferred using iBlot 2 nitrocellulose transfer stacks (Invitrogen). Membranes were blocked for 1Yh and probed with antibodies: APOE 1:1000 (Millipore, AB947), ABCA1 1:1000 (Abcam, 018180), LAMP1 (D2D11) 1:1000 (Cell Signaling Technology 9091S), RAB5 (C8B1) 1:1000 (Cell Signaling Technology, 3547S), ACTIN 1:10,000 (Sigma-Aldrich, A5441) in 5% non-fat dry milk in PBS/0.1% Tween-20 buffer overnight at 4Y°C. Secondary antibody staining 1:10000 was applied for 1Yh at RT, visualized using WesternBright ECL HRP Substrate Kit (Advansta, K-12045), and measured using iBrigh imagining system (Applied Bioscience). Images were analyzed using ImageJ (NIH). Uncropped western blot images are pasted in Figure S7.

### Quantification and statistical analysis

Data were analyzed and visualized in GraphPad Prism 9 (GraphPad Software). In each analysis, three to six independent experiments were performed. Differences of means between groups were tested using a paired t.test All data are represented as group mean ± standard error of the mean (SEM).

## References

1. Bellenguez, C., Küçükali, F., Jansen, I.E., Kleineidam, L., Moreno-Grau, S., Amin, N., Naj, A.C., Campos-Martin, R., Grenier-Boley, B., Andrade, V., et al. (2022). New insights into the genetic etiology of Alzheimer’s disease and related dementias. Nat. Genet. 54, 412–436.

2. Genin, E., Hannequin, D., Wallon, D., Sleegers, K., Hiltunen, M., Combarros, O., Bullido, M.J., Engelborghs, S., De Deyn, P., Berr, C., et al. (2011). APOE and Alzheimer disease: a major gene with semi-dominant inheritance. Mol. Psychiatry 16, 903–907.

3. Belloy, M.E., Andrews, S.J., Le Guen, Y., Cuccaro, M., Farrer, L.A., Napolioni, V., and Greicius, M.D. (2023). APOE Genotype and Alzheimer Disease Risk Across Age, Sex, and Population Ancestry. JAMA Neurol. 10.1001/jamaneurol.2023.3599.

4. Huang, K.-L., Marcora, E., Pimenova, A.A., Di Narzo, A.F., Kapoor, M., Jin, S.C., Harari, O., Bertelsen, S., Fairfax, B.P., Czajkowski, J., et al. (2017). A common haplotype lowers PU.1 expression in myeloid cells and delays onset of Alzheimer’s disease. Nat. Neurosci. 20, 1052–1061.

5. Nott, A., Holtman, I.R., Coufal, N.G., Schlachetzki, J.C.M., Yu, M., Hu, R., Han, C.Z., Pena, M., Xiao, J., Wu, Y., et al. (2019). Brain cell type-specific enhancer-promoter interactome maps and disease risk association. Science. 10.1126/science.aay0793.

6. Novikova, G., Kapoor, M., Tcw, J., Abud, E.M., Efthymiou, A.G., Chen, S.X., Cheng, H., Fullard, J.F., Bendl, J., Liu, Y., et al. (2021). Integration of Alzheimer’s disease genetics and myeloid genomics identifies disease risk regulatory elements and genes. Nat. Commun. 12, 1610.

7. Kosoy, R., Fullard, J.F., Zeng, B., Bendl, J., Dong, P., Rahman, S., Kleopoulos, S.P., Shao, Z., Girdhar, K., Humphrey, J., et al. (2022). Genetics of the human microglia regulome refines Alzheimer’s disease risk loci. Nat. Genet. 54, 1145–1154.

8. Kunkle, B.W., Grenier-Boley, B., Sims, R., Bis, J.C., Damotte, V., Naj, A.C., Boland, A., Vronskaya, M., van der Lee, S.J., Amlie-Wolf, A., et al. (2019). Genetic meta-analysis of diagnosed Alzheimer’s disease identifies new risk loci and implicates Aβ, tau, immunity and lipid processing. Nat. Genet. 51, 414–430.

9. Jones, L., Holmans, P.A., Hamshere, M.L., Harold, D., Moskvina, V., Ivanov, D., Pocklington, A., Abraham, R., Hollingworth, P., Sims, R., et al. (2010). Genetic evidence implicates the immune system and cholesterol metabolism in the aetiology of Alzheimer’s disease. PLoS One 5, e13950.

10. Romero-Molina, C., Garretti, F., Andrews, S.J., Marcora, E., and Goate, A.M. (2022). Microglial efferocytosis: Diving into the Alzheimer’s disease gene pool. Neuron 110, 3513– 3533.

11. Podleśny-Drabiniok, A., Marcora, E., and Goate, A.M. (2020). Microglial Phagocytosis: A Disease-Associated Process Emerging from Alzheimer’s Disease Genetics. Trends Neurosci. 43, 965–979.

12. Andrews, S.J., Renton, A.E., Fulton-Howard, B., Podlesny-Drabiniok, A., Marcora, E., and Goate, A.M. (2023). The complex genetic architecture of Alzheimer’s disease: novel insights and future directions. eBioMedicine 90. 10.1016/j.ebiom.2023.104511.

13. Doran, A.C., Yurdagul, A., Jr, and Tabas, I. (2019). Efferocytosis in health and disease. Nat. Rev. Immunol. 10.1038/s41577-019-0240-6.

14. Deczkowska, A., Keren-Shaul, H., Weiner, A., Colonna, M., Schwartz, M., and Amit, I. (2018). Disease-Associated Microglia: A Universal Immune Sensor of Neurodegeneration. Cell 173, 1073–1081.

15. Keren-Shaul, H., Spinrad, A., Weiner, A., Matcovitch-Natan, O., Dvir-Szternfeld, R., Ulland, T.K., David, E., Baruch, K., Lara-Astaiso, D., Toth, B., et al. (2017). A Unique Microglia Type Associated with Restricting Development of Alzheimer’s Disease. Cell 169, 1276– 1290.e17.

16. Jaitin, D.A., Adlung, L., Thaiss, C.A., Weiner, A., Li, B., Descamps, H., Lundgren, P., Bleriot, C., Liu, Z., Deczkowska, A., et al. (2019). Lipid-Associated Macrophages Control Metabolic Homeostasis in a Trem2-Dependent Manner. Cell 178, 686–698.e14.

17. Krasemann, S., Madore, C., Cialic, R., Baufeld, C., Calcagno, N., El Fatimy, R., Beckers, L., O’Loughlin, E., Xu, Y., Fanek, Z., et al. (2017). The TREM2-APOE Pathway Drives the Transcriptional Phenotype of Dysfunctional Microglia in Neurodegenerative Diseases. Immunity 47, 566–581.e9.

18. Cantoni, C., Bollman, B., Licastro, D., Xie, M., Mikesell, R., Schmidt, R., Yuede, C.M., Galimberti, D., Olivecrona, G., Klein, R.S., et al. (2015). TREM2 regulates microglial cell activation in response to demyelination in vivo. Acta Neuropathol. 129, 429–447.

19. Dolan, M.-J., Therrien, M., Jereb, S., Kamath, T., Gazestani, V., Atkeson, T., Marsh, S.E., Goeva, A., Lojek, N.M., Murphy, S., et al. (2023). Exposure of iPSC-derived human microglia to brain substrates enables the generation and manipulation of diverse transcriptional states in vitro. Nat. Immunol. 24, 1382–1390.

20. Podleśny-Drabiniok, A., Novikova, G., Liu, Y., Dunst, J., Temizer, R., Giannarelli, C., Marro, S., Kreslavsky, T., Marcora, E., and Goate, A.M. (2024). BHLHE40/41 regulate microglia and peripheral macrophage responses associated with Alzheimer’s disease and other disorders of lipid-rich tissues. Nat. Commun. 15, 2058.

21. Cantuti-Castelvetri, L., Fitzner, D., Bosch-Queralt, M., Weil, M.-T., Su, M., Sen, P., Ruhwedel, T., Mitkovski, M., Trendelenburg, G., Lütjohann, D., et al. (2018). Defective cholesterol clearance limits remyelination in the aged central nervous system. Preprint, 10.1126/science.aan4183 10.1126/science.aan4183.

22. Wang, N., Wang, M., Jeevaratnam, S., Rosenberg, C., Ikezu, T.C., Shue, F., Doss, S.V., Alnobani, A., Martens, Y.A., Wren, M., et al. (2022). Opposing effects of apoE2 and apoE4 on microglial activation and lipid metabolism in response to demyelination. Mol. Neurodegener. 17, 75.

23. Poliani, P.L., Wang, Y., Fontana, E., Robinette, M.L., Yamanishi, Y., Gilfillan, S., and Colonna, M. (2015). TREM2 sustains microglial expansion during aging and response to demyelination. J. Clin. Invest. 125, 2161–2170.

24. Nugent, A.A., Lin, K., Van Lengerich, B., Lianoglou, S., Przybyla, L., Davis, S.S., Llapashtica, C., Wang, J., Kim, D.J., Xia, D., et al. (2019). TREM2 Regulates Microglial Cholesterol Metabolism Upon Chronic Phagocytic Challenge. 10.2139/ssrn.3444596.

25. McQuade, A., Kang, Y.J., Hasselmann, J., Jairaman, A., Sotelo, A., Coburn, M., Shabestari, S.K., Chadarevian, J.P., Fote, G., Tu, C.H., et al. (2020). Gene expression and functional deficits underlie TREM2-knockout microglia responses in human models of Alzheimer’s disease. Nat. Commun. 11, 5370.

26. Cignarella, F., Filipello, F., Bollman, B., Cantoni, C., Locca, A., Mikesell, R., Manis, M., Ibrahim, A., Deng, L., Benitez, B.A., et al. (2020). TREM2 activation on microglia promotes myelin debris clearance and remyelination in a model of multiple sclerosis. Acta Neuropathol. 140, 513–534.

27. Bosco, D.B., Kremen, V., Haruwaka, K., Zhao, S., Wang, L., Ebner, B.A., Zheng, J., Xie, M., Dheer, A., Perry, J.F., et al. (2024). Microglial TREM2 promotes phagocytic clearance of damaged neurons after status epilepticus. Brain Behav. Immun. 10.1016/j.bbi.2024.09.034.

28. Bouchon, A., Hernández-Munain, C., Cella, M., and Colonna, M. (2001). A DAP12-mediated pathway regulates expression of CC chemokine receptor 7 and maturation of human dendritic cells. J. Exp. Med. 194, 1111–1122.

29. Turnbull, I.R., Gilfillan, S., Cella, M., Aoshi, T., Miller, M., Piccio, L., Hernandez, M., and Colonna, M. (2006). Cutting edge: TREM-2 attenuates macrophage activation. J. Immunol. 177, 3520–3524.

30. Rangaraju, S., Dammer, E.B., Raza, S.A., Rathakrishnan, P., Xiao, H., Gao, T., Duong, D.M., Pennington, M.W., Lah, J.J., Seyfried, N.T., et al. (2018). Identification and therapeutic modulation of a pro-inflammatory subset of disease-associated-microglia in Alzheimer’s disease. Mol. Neurodegener. 13, 24.

31. Abud, E.M., Ramirez, R.N., Martinez, E.S., Healy, L.M., Nguyen, C.H.H., Newman, S.A., Yeromin, A.V., Scarfone, V.M., Marsh, S.E., Fimbres, C., et al. (2017). iPSC-Derived Human Microglia-like Cells to Study Neurological Diseases. Neuron 94, 278–293.e9.

32. McQuade, A., Coburn, M., Tu, C.H., Hasselmann, J., Davtyan, H., and Blurton-Jones, M. (2018). Development and validation of a simplified method to generate human microglia from pluripotent stem cells. Mol. Neurodegener. 13, 67.

33. Martinez, F.O., Helming, L., Milde, R., Varin, A., Melgert, B.N., Draijer, C., Thomas, B., Fabbri, M., Crawshaw, A., Ho, L.P., et al. (2013). Genetic programs expressed in resting and IL-4 alternatively activated mouse and human macrophages: similarities and differences. Blood 121, e57–e69.

34. Ramachandran, P., Dobie, R., Wilson-Kanamori, J.R., Dora, E.F., Henderson, B.E.P., Luu, N.T., Portman, J.R., Matchett, K.P., Brice, M., Marwick, J.A., et al. (2019). Resolving the fibrotic niche of human liver cirrhosis at single-cell level. Nature 575, 512–518.

35. Fernandez, D.M., Rahman, A.H., Fernandez, N.F., Chudnovskiy, A., Amir, E.-A.D., Amadori, L., Khan, N.S., Wong, C.K., Shamailova, R., Hill, C.A., et al. (2019). Single-cell immune landscape of human atherosclerotic plaques. Nat. Med. 25, 1576–1588.

36. Olah, M., Menon, V., Habib, N., Taga, M.F., Ma, Y., Yung, C.J., Cimpean, M., Khairallah, A., Coronas-Samano, G., Sankowski, R., et al. (2020). Single cell RNA sequencing of human microglia uncovers a subset associated with Alzheimer’s disease. Nat. Commun. 11, 6129.

37. Srinivasan, K., Friedman, B.A., Etxeberria, A., Huntley, M.A., van der Brug, M.P., Foreman, O., Paw, J.S., Modrusan, Z., Beach, T.G., Serrano, G.E., et al. (2019). Alzheimer’s patient brain myeloid cells exhibit enhanced aging and unique transcriptional activation. bioRxiv, 610345. 10.1101/610345.

38. Gosselin, D., Skola, D., Coufal, N.G., Holtman, I.R., Schlachetzki, J.C.M., Sajti, E., Jaeger, B.N., O’Connor, C., Fitzpatrick, C., Pasillas, M.P., et al. (2017). An environment-dependent transcriptional network specifies human microglia identity. Science 356. 10.1126/science.aal3222.

39. Yin, Z., Raj, D., Saiepour, N., Van Dam, D., Brouwer, N., Holtman, I.R., Eggen, B.J.L., Möller, T., Tamm, J.A., Abdourahman, A., et al. (2017). Immune hyperreactivity of Aβ plaque-associated microglia in Alzheimer’s disease. Neurobiol. Aging 55, 115–122.

40. Kamphuis, W., Kooijman, L., Schetters, S., Orre, M., and Hol, E.M. (2016). Transcriptional profiling of CD11c-positive microglia accumulating around amyloid plaques in a mouse model for Alzheimer’s disease. Biochim. Biophys. Acta 1862, 1847–1860.

41. Sala Frigerio, C., Wolfs, L., Fattorelli, N., Thrupp, N., Voytyuk, I., Schmidt, I., Mancuso, R., Chen, W.-T., Woodbury, M.E., Srivastava, G., et al. (2019). The Major Risk Factors for Alzheimer’s Disease: Age, Sex, and Genes Modulate the Microglia Response to Aβ Plaques. Cell Rep. 27, 1293–1306.e6.

42. Friedman, B.A., Srinivasan, K., Ayalon, G., Meilandt, W.J., Lin, H., Huntley, M.A., Cao, Y., Lee, S.-H., Haddick, P.C.G., Ngu, H., et al. (2018). Diverse Brain Myeloid Expression Profiles Reveal Distinct Microglial Activation States and Aspects of Alzheimer’s Disease Not Evident in Mouse Models. Cell Rep. 22, 832–847.

43. Hasselmann, J., Coburn, M.A., England, W., Figueroa Velez, D.X., Kiani Shabestari, S., Tu, C.H., McQuade, A., Kolahdouzan, M., Echeverria, K., Claes, C., et al. (2019). Development of a Chimeric Model to Study and Manipulate Human Microglia In Vivo. Neuron 103, 1016– 1033.e10.

44. Gerrits, E., Brouwer, N., Kooistra, S.M., Woodbury, M.E., Vermeiren, Y., Lambourne, M., Mulder, J., Kummer, M., Möller, T., Biber, K., et al. (2021). Distinct amyloid-β and tau-associated microglia profiles in Alzheimer’s disease. Acta Neuropathol. 141, 681–696.

45. Gazestani, V., Kamath, T., Nadaf, N.M., Dougalis, A., Burris, S.J., Rooney, B., Junkkari, A., Vanderburg, C., Pelkonen, A., Gomez-Budia, M., et al. (2023). Early Alzheimer’s disease pathology in human cortex involves transient cell states. Cell 186, 4438–4453.e23.

46. Mathys, H., Davila-Velderrain, J., Peng, Z., Gao, F., Mohammadi, S., Young, J.Z., Menon, M., He, L., Abdurrob, F., Jiang, X., et al. (2019). Single-cell transcriptomic analysis of Alzheimer’s disease. Nature 570, 332–337.

47. Sun, N., Victor, M.B., Park, Y.P., Xiong, X., Scannail, A.N., Leary, N., Prosper, S., Viswanathan, S., Luna, X., Boix, C.A., et al. (2023). Human microglial state dynamics in Alzheimer’s disease progression. Cell 186, 4386–4403.e29.

48. Miedema, A., Gerrits, E., Brouwer, N., Jiang, Q., Kracht, L., Meijer, M., Nutma, E., Peferoen-Baert, R., Pijnacker, A.T.E., Wesseling, E.M., et al. (2022). Brain macrophages acquire distinct transcriptomes in multiple sclerosis lesions and normal appearing white matter. Acta Neuropathol Commun 10, 8.

49. Cochain, C., Vafadarnejad, E., Arampatzi, P., Pelisek, J., Winkels, H., Ley, K., Wolf, D., Saliba, A.-E., and Zernecke, A. (2018). Single-Cell RNA-Seq Reveals the Transcriptional Landscape and Heterogeneity of Aortic Macrophages in Murine Atherosclerosis. Circ. Res. 122, 1661–1674.

50. Major, J., Fletcher, J.E., and Hamilton, T.A. (2002). IL-4 pretreatment selectively enhances cytokine and chemokine production in lipopolysaccharide-stimulated mouse peritoneal macrophages. J. Immunol. 168, 2456–2463.

51. Fort, M.M., Lesley, R., Davidson, N.J., Menon, S., Brombacher, F., Leach, M.W., and Rennick, D.M. (2001). IL-4 exacerbates disease in a Th1 cell transfer model of colitis. The Journal of Immunology 166, 2793–2800.

52. Wightman, D.P., Jansen, I.E., Savage, J.E., Shadrin, A.A., Bahrami, S., Holland, D., Rongve, A., Børte, S., Winsvold, B.S., Drange, O.K., et al. (2022). Author Correction: A genome-wide association study with 1,126,563 individuals identifies new risk loci for Alzheimer’s disease. Nat. Genet. 54, 1062.

53. Jonsson, T., Stefansson, H., Steinberg, S., Jonsdottir, I., Jonsson, P.V., Snaedal, J., Bjornsson, S., Huttenlocher, J., Levey, A.I., Lah, J.J., et al. (2013). Variant of TREM2 associated with the risk of Alzheimer’s disease. N. Engl. J. Med. 368, 107–116.

54. Cheng-Hathaway, P.J., Reed-Geaghan, E.G., Jay, T.R., Casali, B.T., Bemiller, S.M., Puntambekar, S.S., von Saucken, V.E., Williams, R.Y., Karlo, J.C., Moutinho, M., et al. (2018). The Trem2 R47H variant confers loss-of-function-like phenotypes in Alzheimer’s disease. Mol. Neurodegener. 13, 29.

55. Aslam, M.M., Fan, K.-H., Lawrence, E., Bedison, M.A., Snitz, B.E., DeKosky, S.T., Lopez, O.L., Feingold, E., and Kamboh, M.I. (2023). Genome-wide analysis identifies novel loci influencing plasma apolipoprotein E concentration and Alzheimer’s disease risk. Mol Psychiatry 28, 4451–4462.

56. Depuydt, M.A.C., Prange, K.H.M., Slenders, L., Örd, T., Elbersen, D., Boltjes, A., de Jager, S.C.A., Asselbergs, F.W., de Borst, G.J., Aavik, E., et al. (2020). Microanatomy of the Human Atherosclerotic Plaque by Single-Cell Transcriptomics. Circ. Res. 127, 1437–1455.

57. Hemonnot-Girard, A.-L., Meersseman, C., Pastore, M., Garcia, V., Linck, N., Rey, C., Chebbi, A., Jeanneteau, F., Ginsberg, S.D., Lachuer, J., et al. (2022). Comparative analysis of transcriptome remodeling in plaque-associated and plaque-distant microglia during amyloid-β pathology progression in mice. J. Neuroinflammation 19, 234.

58. Phipson, B., Sim, C.B., Porrello, E.R., Hewitt, A.W., Powell, J., and Oshlack, A. (2022). Propeller: Testing for differences in cell type proportions in single cell data. Bioinformatics 38, 4720–4726.

59. Robinson, M.D., McCarthy, D.J., and Smyth, G.K. (2010). edgeR: a Bioconductor package for differential expression analysis of digital gene expression data. Bioinformatics 26, 139– 140.

60. Blanchard, J.W., Akay, L.A., Davila-Velderrain, J., von Maydell, D., Mathys, H., Davidson, S.M., Effenberger, A., Chen, C.-Y., Maner-Smith, K., Hajjar, I., et al. (2022). APOE4 impairs myelination via cholesterol dysregulation in oligodendrocytes. Nature. 10.1038/s41586-022-05439-w.

61. Victor, M.B., Leary, N., Luna, X., Meharena, H.S., Scannail, A.N., Bozzelli, P.L., Samaan, G., Murdock, M.H., von Maydell, D., Effenberger, A.H., et al. (2022). Lipid accumulation induced by APOE4 impairs microglial surveillance of neuronal-network activity. Cell Stem Cell 29, 1197–1212.e8.

62. Marschallinger, J., Iram, T., Zardeneta, M., Lee, S.E., Lehallier, B., Haney, M.S., Pluvinage, J.V., Mathur, V., Hahn, O., Morgens, D.W., et al. (2020). Lipid-droplet-accumulating microglia represent a dysfunctional and proinflammatory state in the aging brain. Nat. Neurosci. 23, 194–208.

63. Haney, M.S., Pálovics, R., Munson, C.N., Long, C., Johansson, P.K., Yip, O., Dong, W., Rawat, E., West, E., Schlachetzki, J.C.M., et al. (2024). APOE4/4 is linked to damaging lipid droplets in Alzheimer’s disease microglia. Nature. 10.1038/s41586-024-07185-7.

64. Claes, C., Danhash, E.P., Hasselmann, J., Chadarevian, J.P., Shabestari, S.K., England, W.E., Lim, T.E., Hidalgo, J.L.S., Spitale, R.C., Davtyan, H., et al. (2021). Plaque-associated human microglia accumulate lipid droplets in a chimeric model of Alzheimer’s disease. Mol. Neurodegener. 16, 50.

65. Wezel, A., Lagraauw, H.M., van der Velden, D., de Jager, S.C.A., Quax, P.H.A., Kuiper, J., and Bot, I. (2015). Mast cells mediate neutrophil recruitment during atherosclerotic plaque progression. Atherosclerosis 241, 289–296.

66. Khaw, Y.M., Cunningham, C., Tierney, A., Sivaguru, M., and Inoue, M. (2020). Neutrophil-selective deletion of Cxcr2 protects against CNS neurodegeneration in a mouse model of multiple sclerosis. J. Neuroinflammation 17, 49.

67. Haage, V., Tuddenham, J.F., Comandante-Lou, N., Bautista, A., Monzel, A., Chiu, R., Fujita, M., Garcia, F.G., Bhattarai, P., Patel, R., et al. (2024). A pharmacological toolkit for human microglia identifies Topoisomerase I inhibitors as immunomodulators for Alzheimer’s disease. bioRxiv. 10.1101/2024.02.06.579103.

68. Aubert, A., Stüder, F., Colombo, B.M., and Mendoza-Parra, M.A. (2021). A core transcription regulatory circuitry defining microglia cell identity inferred from the reanalysis of multiple human microglia differentiation protocols. Brain Sci. 11, 1338.

69. Pimenova, A.A., Herbinet, M., Gupta, I., Machlovi, S.I., Bowles, K.R., Marcora, E., and Goate, A.M. (2021). Alzheimer’s-associated PU.1 expression levels regulate microglial inflammatory response. Neurobiol. Dis. 148, 105217.

70. Colombo, A., Dinkel, L., Müller, S.A., Sebastian Monasor, L., Schifferer, M., Cantuti-Castelvetri, L., König, J., Vidatic, L., Bremova-Ertl, T., Lieberman, A.P., et al. (2021). Loss of NPC1 enhances phagocytic uptake and impairs lipid trafficking in microglia. Nat. Commun. 12, 1158.

71. Orihuela, R., McPherson, C.A., and Harry, G.J. (2016). Microglial M1/M2 polarization and metabolic states. Br. J. Pharmacol. 173, 649–665.

72. Sangineto, M., Ciarnelli, M., Cassano, T., Radesco, A., Moola, A., Bukke, V.N., Romano, A., Villani, R., Kanwal, H., Capitanio, N., et al. (2023). Metabolic reprogramming in inflammatory microglia indicates a potential way of targeting inflammation in Alzheimer’s disease. Redox Biol 66, 102846.

73. Pearce, E.L., and Pearce, E.J. (2013). Metabolic pathways in immune cell activation and quiescence. Immunity 38, 633–643.

74. Lee, S., Devanney, N.A., Golden, L.R., Smith, C.T., Schwartz, J.L., Walsh, A.E., Clarke, H.A., Goulding, D.S., Allenger, E.J., Morillo-Segovia, G., et al. (2023). APOE modulates microglial immunometabolism in response to age, amyloid pathology, and inflammatory challenge. Cell Rep. 42, 112196.

75. March-Diaz, R., Lara-Ureña, N., Romero-Molina, C., Heras-Garvin, A., Luis, C.O.S., Alvarez-Vergara, M.I., Sanchez-Garcia, M.A., Sanchez-Mejias, E., Davila, J.C., Rosales-Nieves, A.E., et al. (2021). Hypoxia compromises the mitochondrial metabolism of Alzheimer’s disease microglia via HIF1. Nat Aging 1, 385–399.

76. Ulland, T.K., Song, W.M., Huang, S.C.-C., Ulrich, J.D., Sergushichev, A., Beatty, W.L., Loboda, A.A., Zhou, Y., Cairns, N.J., Kambal, A., et al. (2017). TREM2 Maintains Microglial Metabolic Fitness in Alzheimer’s Disease. Cell 170, 649–663.e13.

77. Castoldi, A., Monteiro, L.B., van Teijlingen Bakker, N., Sanin, D.E., Rana, N., Corrado, M., Cameron, A.M., Hässler, F., Matsushita, M., Caputa, G., et al. (2020). Triacylglycerol synthesis enhances macrophage inflammatory function. Nat. Commun. 11, 4107.

78. Prakash, P., Manchanda, P., Paouri, E., Bisht, K., Sharma, K., Wijewardhane, P.R., Randolph, C.E., Clark, M.G., Fine, J., Thayer, E.A., et al. (2023). Amyloid β Induces Lipid Droplet-Mediated Microglial Dysfunction in Alzheimer’s Disease. bioRxiv. 10.1101/2023.06.04.543525.

79. Breland, U.M., Halvorsen, B., Hol, J., Øie, E., Paulsson-Berne, G., Yndestad, A., Smith, C., Otterdal, K., Hedin, U., Waehre, T., et al. (2008). A potential role of the CXC chemokine GROalpha in atherosclerosis and plaque destabilization: downregulatory effects of statins. Arterioscler. Thromb. Vasc. Biol. 28, 1005–1011.

80. Yin, Z., Rosenzweig, N., Kleemann, K.L., Zhang, X., Brandão, W., Margeta, M.A., Schroeder, C., Sivanathan, K.N., Silveira, S., Gauthier, C., et al. (2023). APOE4 impairs the microglial response in Alzheimer’s disease by inducing TGFβ-mediated checkpoints. Springer Nature, 1839–1853.

81. Safaiyan, S., Kannaiyan, N., Snaidero, N., Brioschi, S., Biber, K., Yona, S., Edinger, A.L., Jung, S., Rossner, M.J., and Simons, M. (2016). Age-related myelin degradation burdens the clearance function of microglia during aging. Nat. Neurosci. 19, 995–998.

82. Butler, A., Hoffman, P., Smibert, P., Papalexi, E., and Satija, R. (2018). Integrating single-cell transcriptomic data across different conditions, technologies, and species. Nat Biotechnol 36, 411–420.

83. Korsunsky, I., Millard, N., Fan, J., Slowikowski, K., Zhang, F., Wei, K., Baglaenko, Y., Brenner, M., Loh, P.-R., and Raychaudhuri, S. (2019). Fast, sensitive and accurate integration of single-cell data with Harmony. Nat. Methods 16, 1289–1296.

84. Amezquita, R.A., Lun, A.T.L., Becht, E., Carey, V.J., Carpp, L.N., Geistlinger, L., Marini, F., Rue-Albrecht, K., Risso, D., Soneson, C., et al. (2020). Publisher Correction: Orchestrating single-cell analysis with Bioconductor. Nat Methods 17, 242.

85. Kiselev, V.Y., Yiu, A., and Hemberg, M. (2018). scmap: projection of single-cell RNA-seq data across data sets. Nat. Methods 15, 359–362.

86. Hoffman, G.E., and Roussos, P. (2021). Dream: powerful differential expression analysis for repeated measures designs. Bioinformatics 37, 192–201.

87. Cahill, K.M., Huo, Z., Tseng, G.C., Logan, R.W., and Seney, M.L. (2018). Improved identification of concordant and discordant gene expression signatures using an updated rank-rank hypergeometric overlap approach. Sci. Rep. 8, 9588.

88. Mohamed, A., Molendijk, J., and Hill, M.M. (2020). lipidr: A Software Tool for Data Mining and Analysis of Lipidomics Datasets. J. Proteome Res. 19, 2890–2897.

89. Machlovi, S.I., Neuner, S.M., Hemmer, B.M., Khan, R., Liu, Y., Huang, M., Zhu, J.D., Castellano, J.M., Cai, D., Marcora, E., et al. (2022). APOE4 confers transcriptomic and functional alterations to primary mouse microglia. Neurobiol. Dis. 164, 105615.

90. Kolberg, L., Raudvere, U., Kuzmin, I., Adler, P., Vilo, J., and Peterson, H. (2023). g:Profiler-interoperable web service for functional enrichment analysis and gene identifier mapping (2023 update). Nucleic Acids Res. 51, W207–W212.

91. Subramanian, A., Tamayo, P., Mootha, V.K., Mukherjee, S., Ebert, B.L., Gillette, M.A., Paulovich, A., Pomeroy, S.L., Golub, T.R., Lander, E.S., et al. (2005). Gene set enrichment analysis: a knowledge-based approach for interpreting genome-wide expression profiles. Proc. Natl. Acad. Sci. U. S. A. 102, 15545–15550.

92. Hao, Y., Stuart, T., Kowalski, M.H., Choudhary, S., Hoffman, P., Hartman, A., Srivastava, A., Molla, G., Madad, S., Fernandez-Granda, C., et al. (2024). Dictionary learning for integrative, multimodal and scalable single-cell analysis. Nat. Biotechnol. 42, 293–304.

93. Hafemeister, C., and Satija, R. (2019). Normalization and variance stabilization of single-cell RNA-seq data using regularized negative binomial regression. Genome Biol. 20, 296.

