## Supplementary Figures for "Cytokine-induced reprogramming of human macrophages toward Alzheimer’s disease-relevant molecular and cellular phenotypes *in vitro*"

### Supplementary figures and supplementary tables legends

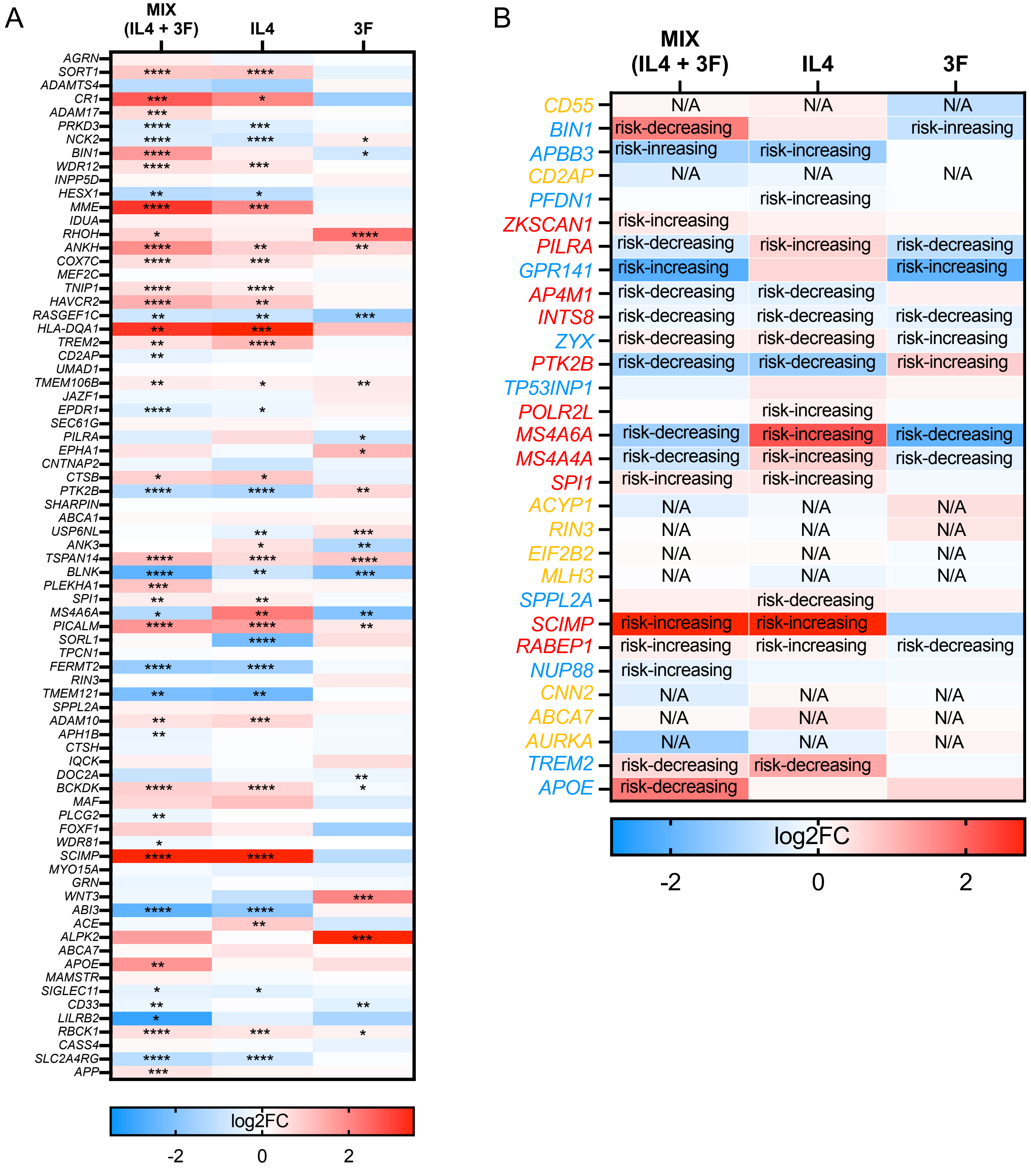


**Figure S1. Cytokine treatment altered the expression level of candidate AD risk genes.** A) Heatmap showing the effect sizes of genes prioritized in 81 AD/dementia genome-wide significant loci by Bellenguez et al., 2022 and Wightman et al., 2022 in cytokine mix, IL4, and 3F treated macrophages as compared to control. B) Heatmap showing the effect sizes of myeloid candidate causal genes nominated through SMR and Hi-C approaches by Novikova et al., 2021. Red indicates that increased expression is associated with increased AD risk; blue indicates that decreased expression is associated with increased AD risk; gold indicates that risk association cannot be reliably inferred and was marked as “N/A”. “risk-decreasing” suggests the expression of an AD risk gene in cytokine-treated conditions is modulated in the opposite direction to the association with AD risk. “risk-increasing” indicates the expression of an AD risk gene in cytokine-treated conditions is modulated in the same direction as the association with AD risk found by Novikova et al., 2021. Empty cells indicate that certain genes with known AD risk associations were not differentially expressed (adj.p > 0.05).

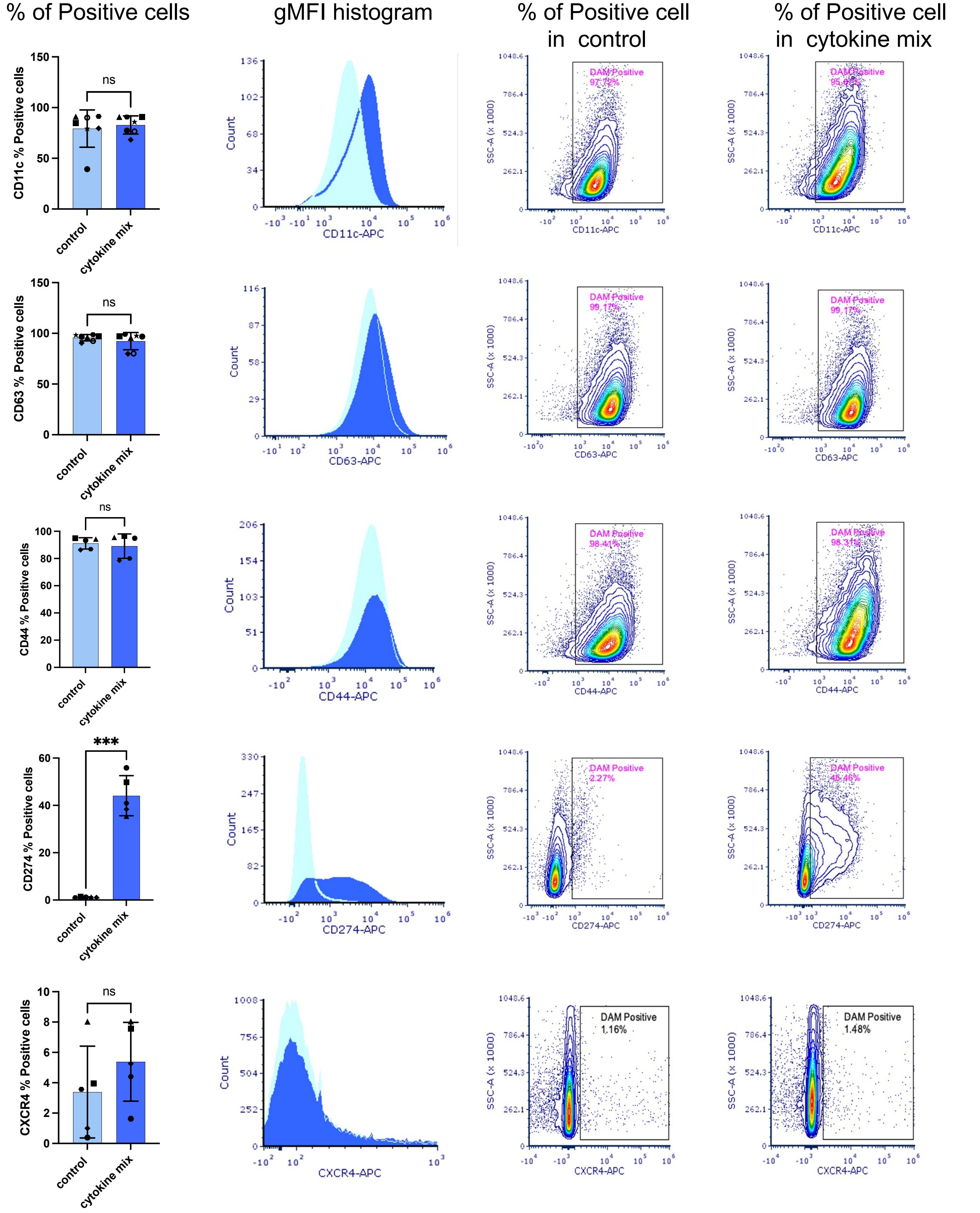


**Figure S2. Quantification of surface DLAM markers by flow cytometry reveals no change in the percentage of positive cells (corresponding to Figure 2C).** Quantification of percentage of positive cells (based on fluorescent-minus-one control). Quantification on the left, representative histograms and dot plots on the right. Groups were tested by paired t.test (two-tailed) * $p < 0.05$, ** $p < 0.01$, *** $p < 0.001$, ns – not significant

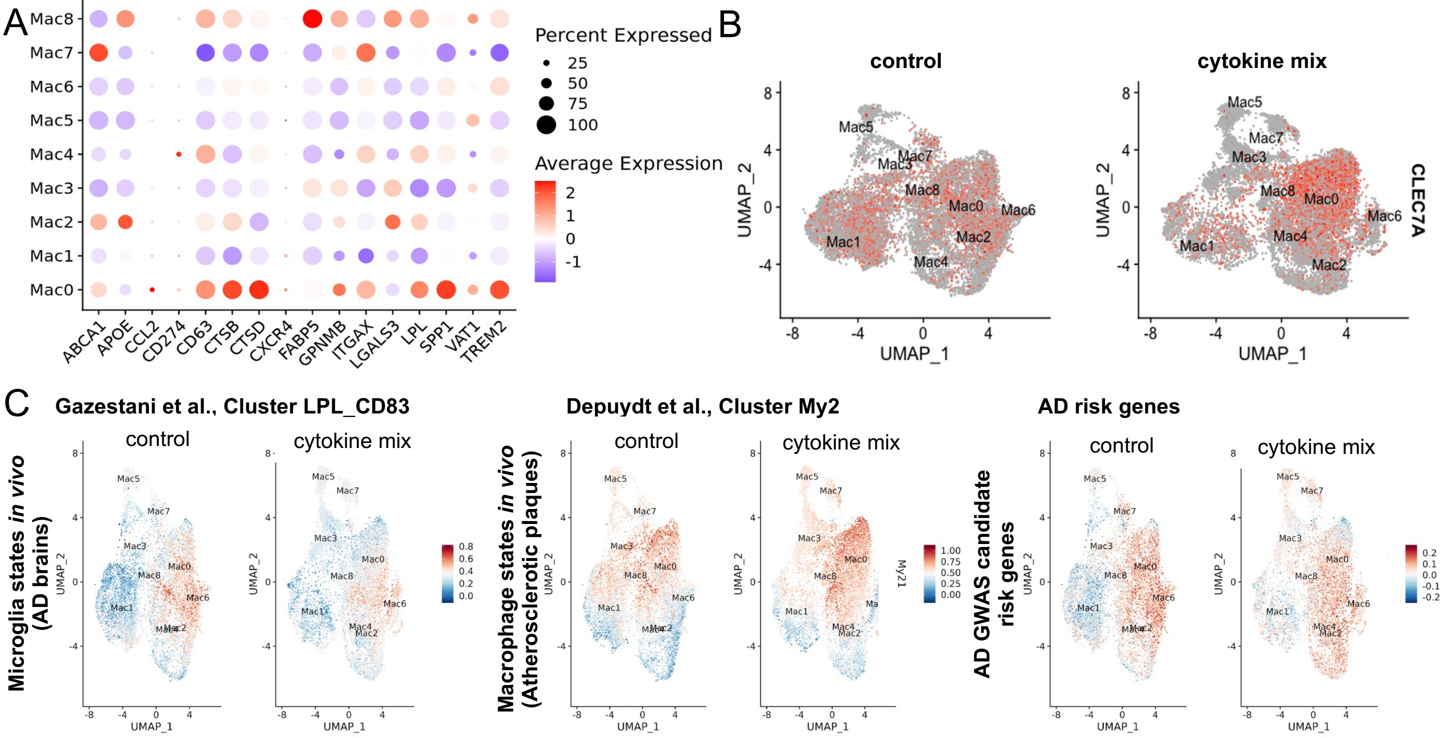


D

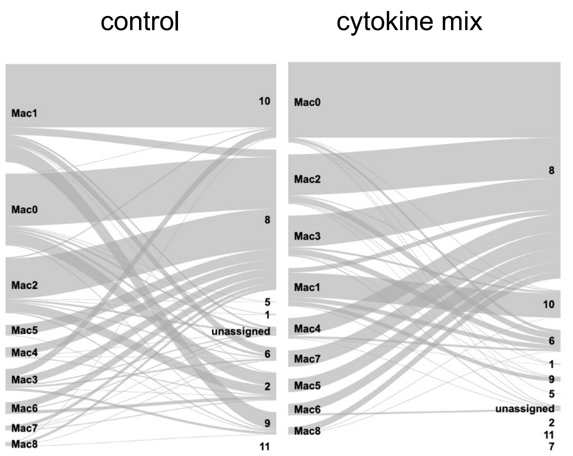


Reference clusters from Dolan et al, 2023

| Query clusters from this study | DAM Cl. 2 | DAM Cl. 8 | Prolif Cl.10 | Proli Cl.6 | Prolif Cl.9 | DAM Cl. 2 | DAM Cl. 8 | Prolif Cl.10 | Proli Cl.6 | Prolif Cl.9 |
|---|---|---|---|---|---|---|---|---|---|---|
| Mac0 | 601 | 2,476 | | 117 | | | 3,736 | | 147 | |
| Mac1 | | 356 | 3,019 | 356 | 806 | | 235 | 1,199 | 218 | 202 |
| Mac2 | 443 | 1,931 | | | | | 2,016 | | 304 | |
| Mac3 | | 391 | 381 | | 115 | | 1,547 | | 290 | |
| Mac4 | | 397 | | | | | 884 | | | |
| Mac5 | | 490 | | | | | 657 | | | |
| Mac6 | 154 | 349 | | 117 | 115 | | 474 | | | |
| Mac7 | | 167 | | | | | 813 | | | |
| Mac8 | | 128 | | | | | 341 | | | |
| Total | 1,198 | 6,685 | 3,400 | 590 | 1,036 | 0 | 10,703 | 1,199 | 959 | 202 |

control (n = 13,022 cells) | cytokine mix (n = 13,063 cells)

**Figure S3. Expression of DLAM genes and transcriptional programs in THP-1 macrophages treated with cytokine mix reveals increased proportion of DLAM cells** A) Dotplot showing expression of selected DLAM marker genes across all clusters. B) Expression of CLEC7A across clusters and split by treatment condition. Higher expression shown in red and low to no expression in grey. C) UMAP projection of THP-1 macrophages dataset, cells colored by module scores of transcriptional signatures identified DAM cluster LPL_CD84 in AD biopsies identified by Gazestani et al., 2024; LAM cluster My2 of ABCG1$^{+}$ foamy macrophages from human atherosclerotic plaques reported by Depuydt et al, 2020, and AD risk genes identified in GWAS study and nominated and candidate causal genes in GWAS loci (Bellenguez et al, 2022) D) Sankey diagrams showing the scmap-cluster projection of query clusters (this study) to reference clusters (Dolan et al, 2023) in control (No cytokines) and treatment (Cytokines) conditions. The number of cells projecting from one cluster to another is listed in a table next to Sankey's diagrams. Projection < 100 cells were not listed. In ”No cytokines” condition 103 cells projected to “unassigned” cluster which, for clarity, is not listed in the table.

A

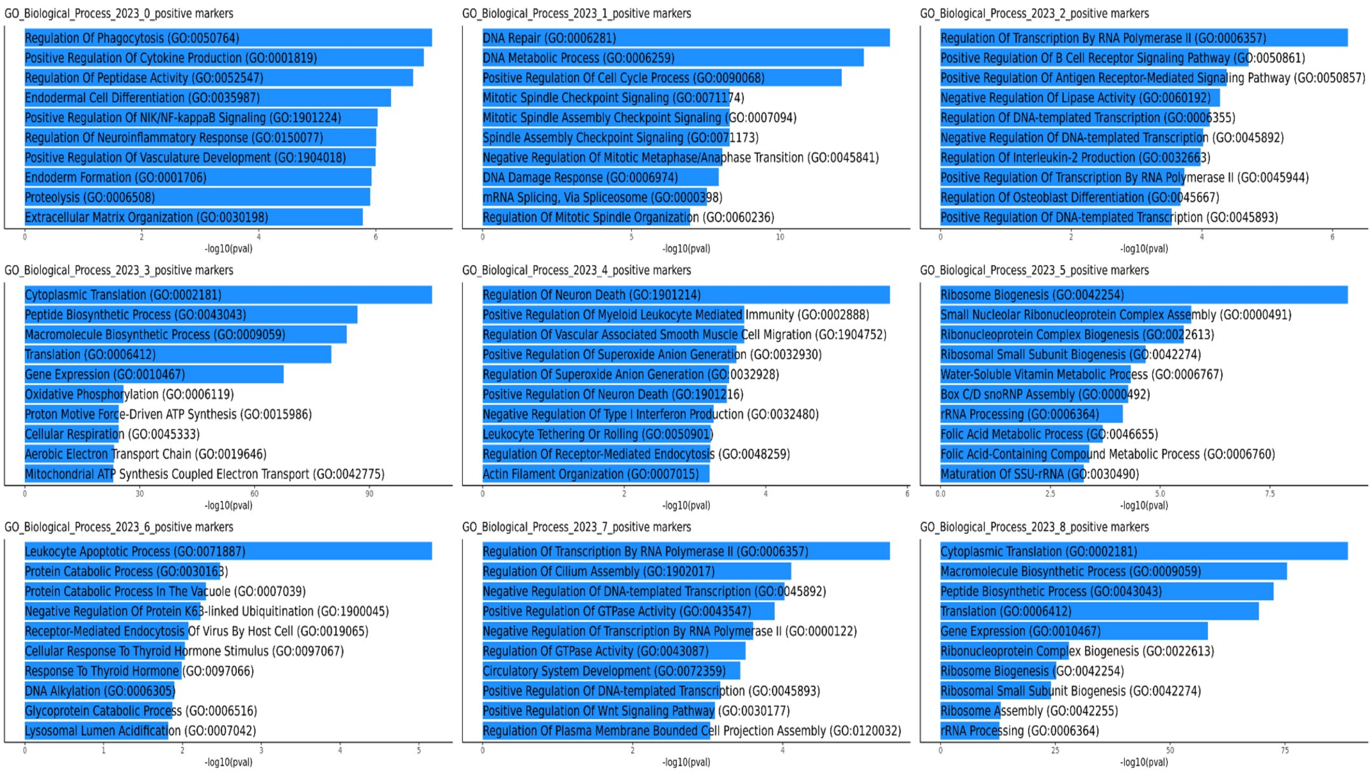


B

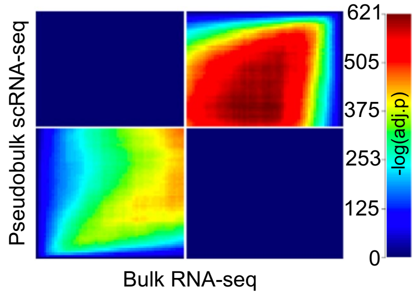


**Figure S4 Enrichment of gene ontology terms in scRNAseq clusters and comparison of transcriptome identified in scRNAseq and bulk RNAseq** A) Enrichment of biological pathways in single cell clusters used for cluster annotation B) Rank rank hypergeometric overlap of bulk RNA-seq vs pseudobulk scRNA-seq differentially expressed genes between treatment conditions

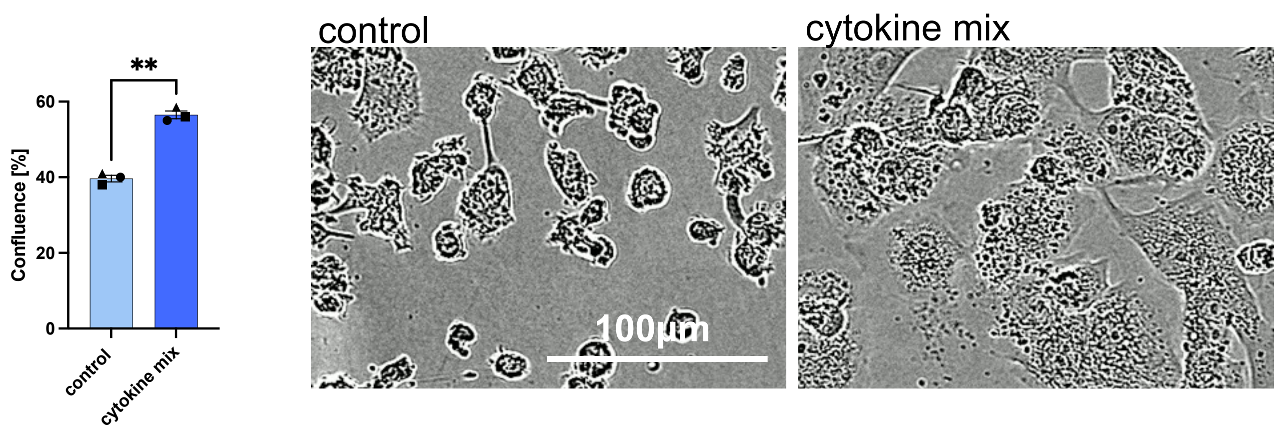


**Figure S5: Surface area of cytokine mix-treated THP-1 macrophages is increased** A) Quantification of area occupied by macrophages (left); representative images (right). Groups were tested by t.test. ** $p < 0.01$

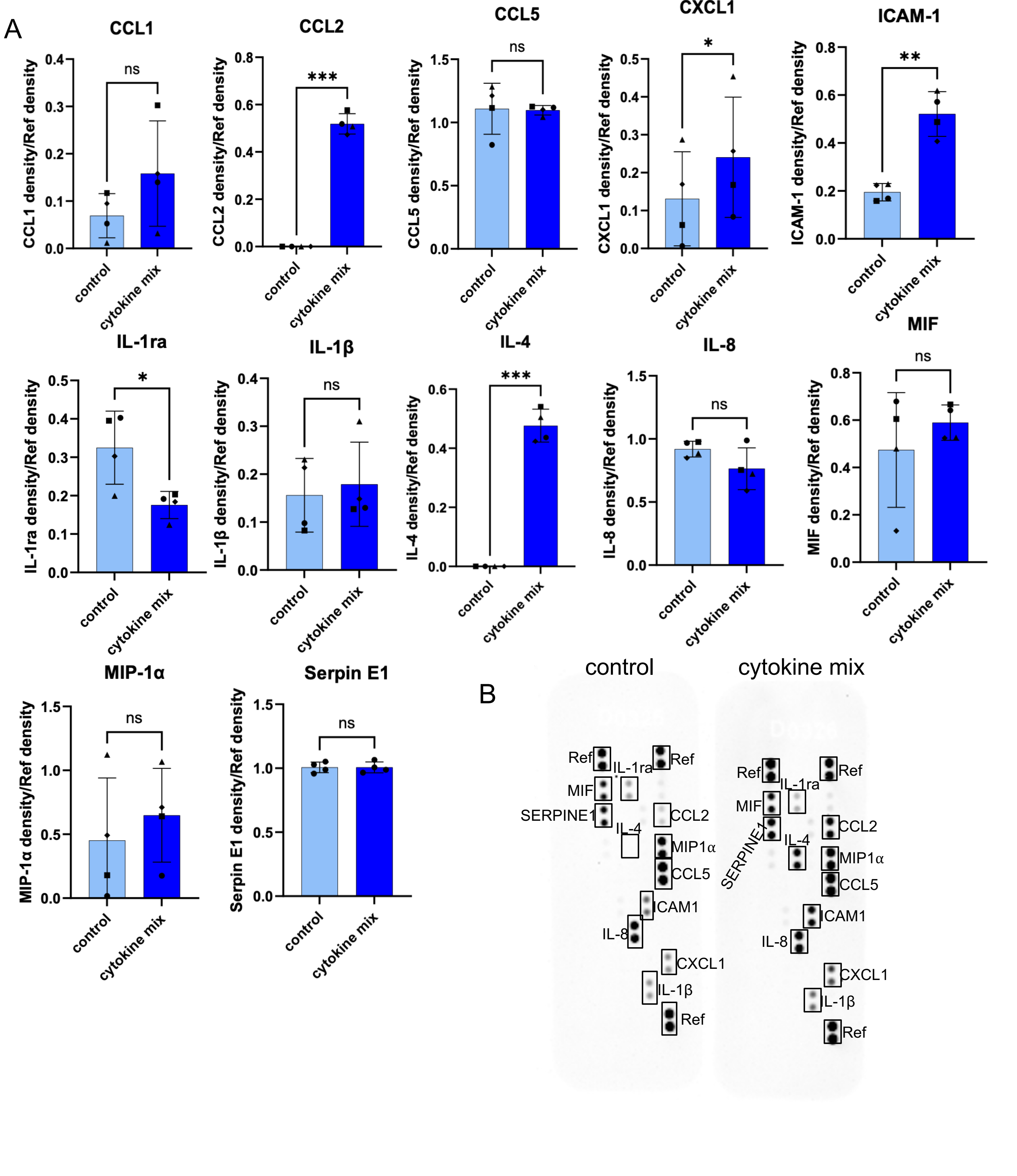


**Figure S6: Secretion of pro- and anti-inflammatory markers in cytokine-treated THP1 macrophages reveal; changes in DLAM cytokines and adhesive molecules** A) Quantifications and dot blots of cytokines secreted in conditioned media from THP-1 macrophages treated with cytokines measured using the Proteome Profiler Human Cytokine Array. Each symbol represents independent macrophage differentiation. Groups were tested by paired t.test (two-tailed) * $p < 0.05$, ** $p < 0.01$, *** $p < 0.001$, ns – not significant B) Representative dot blots

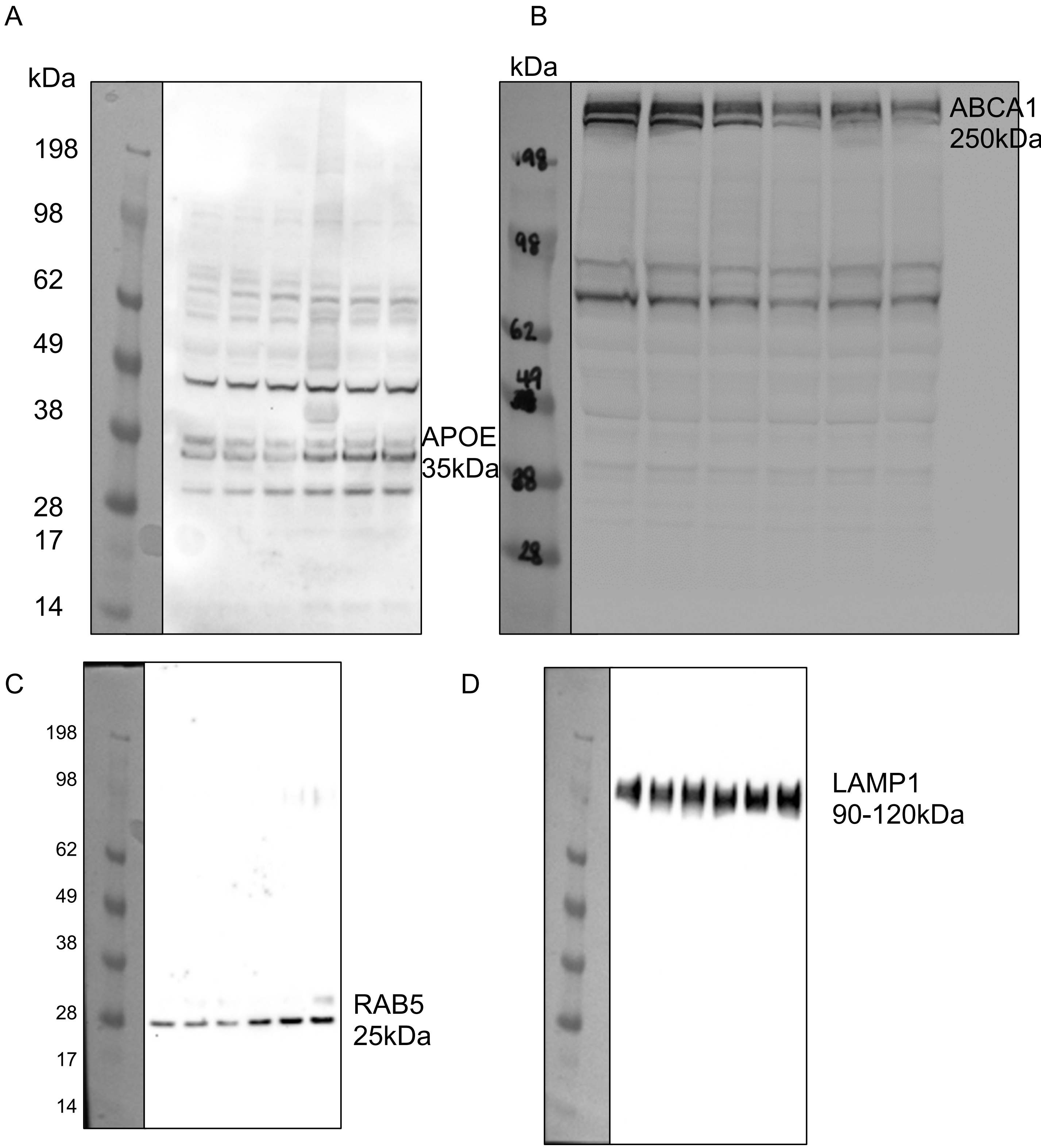


**Figure S7: Uncropped western blot images.** A) APOE expression in cytokine-treated THP-1 macrophages (Figure 6D) B) ABCA1 expression in cytokine-treated THP-1 macrophages (Figure 6F) C-D) RAB5 and LAM1 expression in cytokine mix-treated THP-1 macrophages (Figure 4E3).

#### List of Supplementary Tables

**Table S1: Differential gene expression (DGEA) and gene set enrichment (GSEA) analysis of cytokine-treated and control THP-1 macrophages.**

TMP sheet. Normalized counts in all samples (bulk RNAseq).

DGEA sheet. Differential gene expression analysis between cytokine-treated THP-1 macrophages and control (bulk RNAseq)

GSEA sheet. Gene set enrichment analysis of DGEA.

**Table S2: DLAM, Proliferative, Interferon- and LPS-related gene sets from single cell or single nuclei RNAseq of AD human brains and mouse models**

**DLAM_genesets sheet.** Human and humanized DLAM genesets used in gene sets enrichment analysis

**AD_risk_genes sheet**. Differential expression of prioritized genes in 81 AD risk loci in cytokine-treated THP-1 macrophages. Genes were prioritized by Bellenguez at al, 2022 and Wightman et al, 2022 and listed in Figure 2 by Andrews et al, 2023.

**myeloid_AD_candidate_genes sheet**. Differential expression of myeloid candidate AD risk genes nominated in AD GWAS loci by Novikova et al, 2021. in cytokine-treated THP-1 macrophages. This table corresponds to Supplementary Figure 1B

**Table S3: scRNAseq**

**Cluster_annotation sheet**. Number and percentage of cells per sample, treatment condition and cluster. Cluster annotation is based on enrichment of gene expression signatures from the literature and biological pathways.

**Cluster_marker_genes sheet**. Expression of top marker genes per cluster (log2FC > 0.6 & pct.1 > 0.7 & adj.p < 0.05).

**Cluster_proportion_statistics sheet**. Differences in proportion of cells within each cluster by treatment group statistics using speckle.

**Hypergeometric_overlap sheet**. Raw data for Supplementary Figure 3A. Overlap between published cluster marker genes (gene sets) and cluster marker genes from this study. Gene sets have unique gene sets (GS) identifier and are listed in Table S2

**Pseudobulk_DEG_results_Cytokine_vs_control sheet**. Differential gene expression results of pseudobulked cells between cytokine mix treatment and control conditions using edgeR. [logFC = log fold change of NT vs MIX expression; logCPM = log counts per million; F = ; P-value = p-value of logFC; FDR = FDR-adjusted p-value]

**Table S4: Differential lipid abundance**

**Raw_data sheet**. Raw data listing peak area for each lipid species detected in lipidomic experiment

**Differential_abundance sheet:** Differentially abundant lipid species in control and cytokine-treated THP-1 macrophages. logFC and adj.P-values were obtained using lipidR package
